# Disentangling the sucrose metabolism of the corn smut *Ustilago maydis* reveals unexpected complexity

**DOI:** 10.64898/2026.06.11.731450

**Authors:** Tom Berwanger, Ali Khoshouei, Heba Yehia, Joana Charlot Hasenklever, Tanvir Hassan, Anna Barbara Matuszyńska, Jörg Kämper, Khaled A. Selim, Florian Altegoer, Kerstin Schipper

## Abstract

The race for carbohydrates shapes organismic interactions. In plant pathogenic fungi, sucrose is a key nutrient as it constitutes the major transport sugar in plants. Here, we investigate sucrose acquisition in the corn smut fungus *Ustilago maydis*, a biotrophic pathogen that transitions from yeast-like to hyphal growth for infection. We establish that the fungus encodes a secreted acidic invertase, Suc2, with a dimeric canonical glycoside hydrolase 32 architecture. Comparative biochemical analyses across fungal homologs indicate that this dimeric architecture represents the predominant state, whereas higher-order oligomers, as initially described for *Saccharomyces cerevisiae* Suc2, are restricted to a subset of lineages. Unexpectedly, elimination of Suc2 did not impair yeast-like growth on sucrose. Similarly, deletion of genes for sucrose transporter Srt1 and cytosolic hydrolase Suc1, typically associated with intracellular sucrose metabolism, did not abolish growth. Instead, sucrose utilization during yeast-like growth depended on a repurposed non-canonical module comprising maltose transporter Agt1 and intracellular (iso)maltases. In contrast, pathogenic development strongly relied on the canonical intracellular sucrose utilization pathway mediated by Srt1 and Agt1. Infection was strongly diminished in strains unable to metabolise sucrose, confirming its central nutritional role for the fungus. Together, our work defines the complete sucrose utilization repertoire of *U. maydis* and uncovers a lifestyle-dependent metabolic switch between alternative sucrose acquisition strategies. Flexible carbon acquisition might represent a widespread adaptive strategy in basidiomycete pathogenic fungi.

## 1 Introduction

Sucrose constitutes a ubiquitously abundant carbon source primarily synthesized by photosynthetic organisms (1, 2) and constitutes the major transport form of photoassimilates in plants (3). Many microorganisms, including plant symbionts and pathogens, exploit this disaccharide for feeding (4). Secreted invertases hydrolyze sucrose into the hexoses fructose and glucose for uptake and metabolisation. Alternatively, sucrose can be imported for intracellular hydrolysis, for instance by sucrose phosphatases or intracellular invertases (2).

Sucrose metabolism has traditionally been studied in *Saccharomyces cerevisiae*, a member of the *Saccharomyces* (= sugar fungus) genus (2). This microbial model preferentially employs secreted glycoside hydrolase 32 (GH32) family invertases (β-fructofuranosidase, EC 3.2.1.26) for sucrose consumption. Certain strains lacking invertases survive on sucrose by employing the maltose transporter Agt1 in conjunction with intracellular maltases and isomaltases that enable uptake and slow growth on sucrose (5, 6). However, several isolates, including laboratory strains like S288C, carry a genomic deletion (*mal2*) of the cognate genes and are hence unable to grow on maltose or use the pathway for alternative sucrose consumption (7).

*S. cerevisiae* strains can comprise up to 9 different variants of sucrose hydrolyzing invertases encoded by the *SUC1-9* genes. The number of active alleles varies greatly between different isolates, but *SUC2* is thought to constitute the ancestral gene which is present in all isolates (8). Notably, this multiplication event is only observed in *S. cerevisiae*, while it is lacking in the close relatives like *S. paradoxus* or *S. arboricola* (9), hence probably indicating evolutionary adaptation effects in response to human domestication of *S. cerevisiae*. All *SUC* genes in *S. cerevisiae* encode dual targeted enzymes (10) where the cognate genes are transcribed in two versions, one transcript containing the complete open reading frame including the signal peptide sequence for enzyme secretion and one truncated transcript resulting in a cytoplasmic version (11). While the short version is expressed constitutively at a low basic level, the formation of the long transcript is regulated by the presence of glucose via carbon repression (11). Hence, when glucose levels are low while sucrose is present, secreted ScSuc2 mediates carbon acquisition by extracellular sucrose hydrolysis and subsequent uptake of released hexoses (2). Interestingly, structural studies of *S. cerevisiae* invertase ScSuc2 suggest that it assembles into higher-order oligomers, preferentially octamers (tetramer of dimers) (12, 13).

Sucrose is also thought to be central to the lifestyle of the corn smut fungus *Ustilago maydis*, a biotrophic pathogen of *Zea mays.* This dimorphic fungus grows as a saprotroph in a haploid yeast form that divides by budding. On the plant surface, two compatible yeast cells recognize each other, mate and form an infectious dikaryotic hypha (14). Using an appressorium-like structure, the hypha penetrates the plant tissue and subsequently proliferates intracellularly. To this end, fungal hyphae invade living plant cells by invaginating the plant plasma membrane, thereby establishing a biotrophic interaction zone (15). Importantly, unlike destroying plant tissue as observed for necrotrophs, the fungus keeps host cells alive and reprograms the plant to accumulate nutrients preferentially at infection sites (16). During proliferation, the fungal hyphae particularly spread along the phloem vasculature. Later, fungal infection causes the formation of large tumors that accumulate sucrose and hexoses (17). A large set of sugar transporters is induced in the pathogenic stage of *U. maydis*, and two of them, the high-affinity sucrose transporter Srt1 and the hexose transceptor Hxt1 were shown to significantly contribute to virulence as deletion mutants of the cognate genes show significantly decreased infection symptoms (18, 19). On the plant side, changes in maize sugar transporters were detected upon fungal invasion, fostering the accumulation of intracellular starch and apoplastic soluble sugars in infected regions. Hence, feeding on apoplastic sucrose and hexoses seems central to efficient fungal infection, likely resulting in an intricate race for carbohydrates in infected plants (16).

Based on the assumption that sucrose is a key carbon source during fungal development, we here studied the sucrose metabolism of *U. maydis* during yeast-like axenic growth and infection. We first dissected the role of UmSuc2, an ortholog to *S. cerevisiae* ScSuc2, revealing insights into its role in the context of a plant pathogen. Moreover, we uncovered an additional, unexpectedly large set of sucrose-hydrolyzing enzymes as well as relevant transporters for sucrose import, mediating a highly flexible, context-dependent sucrose metabolization strategy.

## 2 Results

### 2.1 The dimeric secreted invertase Suc2 mediates extracellular sucrose hydrolysis

Bioinformatic annotation suggested that *U. maydis* contains a single secreted invertase, Suc2 (UMAG_01945, UmSuc2), with a canonical glycosyl hydrolase family 32 domain (GH32) for extracellular sucrose hydrolysis (Fig. 1A). In line with that, we detected extracellular sucrose-hydrolysing activity in supernatants of the *U. maydis* laboratory strain AB33 (Fig. 1B). To verify the predicted invertase activity of UmSuc2, we established *Saccharomyces cerevisiae* BY4741 *SUC2Δ*, a strain unable to import or metabolize sucrose, as a heterologous test system (Fig. S1). Cell extracts of tester strains producing UmSuc2 were able to hydrolyse sucrose, similar to control strains containing ScSuc2 (Fig. 1C). This confirms that UmSuc2 functions as an invertase. Sucrose hydrolysis was also detected in culture supernatants of BY4741 *SUC2Δ* equipped with UmSuc2, confirming the presence of a functional signal peptide for secretion (Fig. 1D).

**Figure 1:**
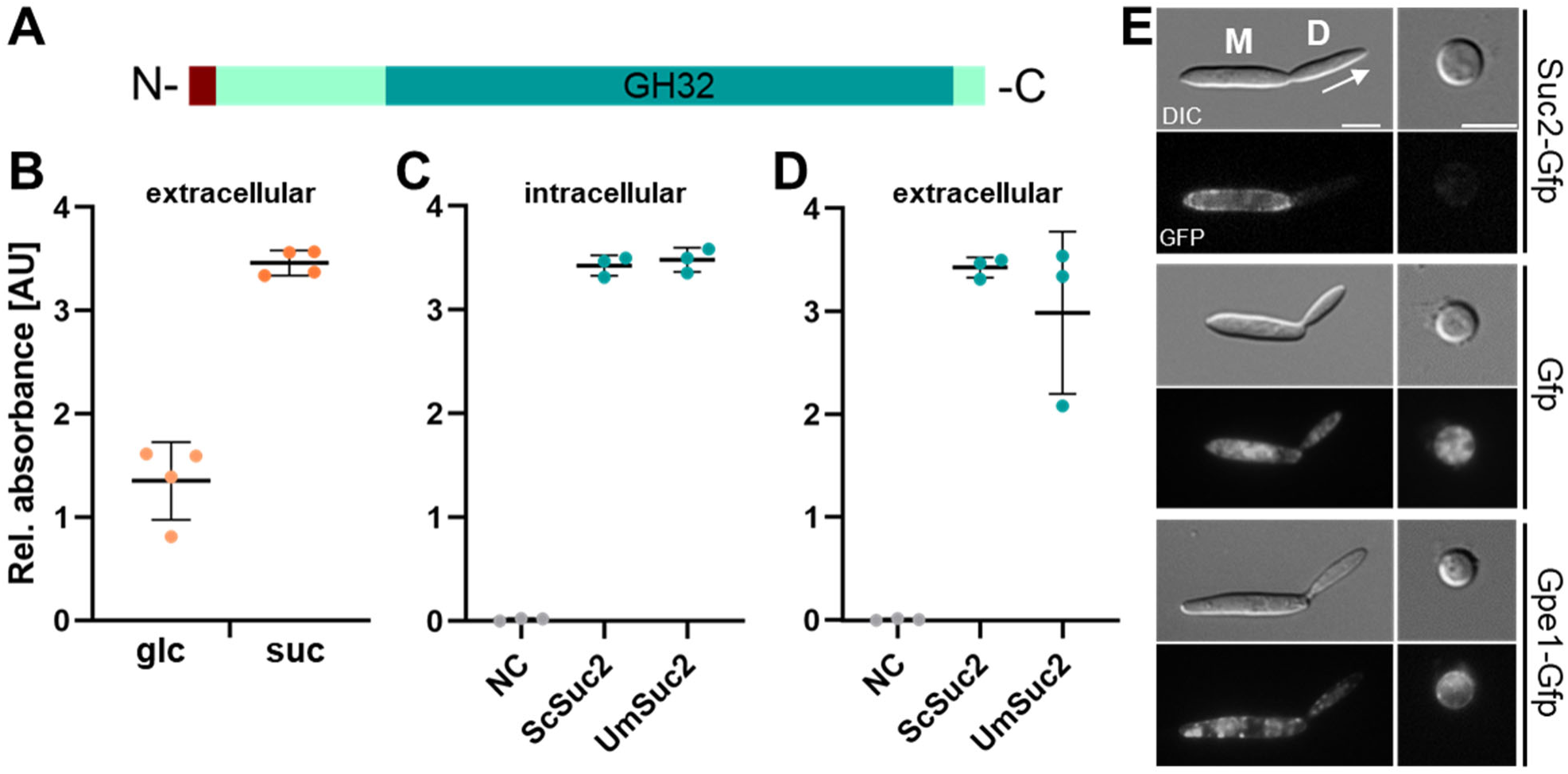
UmSuc2 is a secreted invertase. **A**, Schematic structure of Suc2. Teal, glyco32 domain; brown, predicted N-terminal signal peptide. **B,** DNS assays with sucrose as a substrate using cell-free culture supernatants of *U. maydis* AB33. Cultures were pre-grown on glucose (glc) or sucrose (suc) until the carbon source was depleted. **C-D,** DNS assays with sucrose as a substrate of reporter strain *S. cerevisiae* BY4741 *SUC2Δ* derivatives producing ScSuc2 and UmSuc2. All strains were grown in liquid cultures with glucose until the carbon source was depleted. Then, DNS assays were conducted using cell extracts (**C**) or cell-free culture supernatants (**D**) and sucrose as a substrate to detect reducing sugars resulting from hydrolysis of the disaccharide. Detection of reducing sugars is indicative for sucrose hydrolysis. NC, negative control using BY4741 *SUC2Δ* without additional genes. Assays were conducted in biological triplicates. Error bars depict standard deviation. **E**, Fluorescence microscopic of indicated yeast-like growing strains (left) and protoplasts (right) of *U. maydis*. M, mother cell; D, daughter cell. Scale bar, 5 µm. Two strains are AB33 derivatives producing Suc2-Gfp and Gfp as cytoplasmic control. SG200 producing the G-Protein coupled receptor Gpe1-Gfp (also known as Pit1-Gfp) was used as a control for membrane localization (20, 21).

To visualize the invertase, a genetic construct for UmSuc2 fused to the green fluorescent protein (Suc2-Gfp) was expressed in *U. maydis* AB33. Fluorescence microscopy revealed localization at the cell boundary of yeast-like growing cells, indicating that the protein is secreted and partially trapped at the cell boundary (Fig. 1E, top left panel). Interestingly, Suc2-Gfp only accumulates in the boundary of mother cells, but is absent in daughter cells. In contrast to strains producing intracellular Gfp or the membrane protein Gpe1-Gfp (20, 21), the fluorescent signal disappeared upon removal of the cell wall by lytic enzymes (Fig. 1E, top right panel). This verifies that Suc2-Gfp is indeed exported and found in the cell boundary, but likely not attached to the cell membrane.

To characterize UmSuc2 in more detail, a signal peptide–truncated variant (UmSuc2^ΔSP^) was produced recombinantly in *E. coli* and purified to high homogeneity (Fig. S2A). Biochemical characterization indicates a preference for acidic pH (2-5.5) and a high thermal stability (Fig. S2 B,C). The catalytic activity of UmSuc2^ΔSP^ was evaluated using sucrose as a variable substrate to determine the kinetic constants by fitting initial velocity data to the Michaelis-Menten equation using non-linear regression. Analysis of initial velocity data revealed a K_m_ of 53 mM with a V_max_ of 2.4 mM/min and a turnover K_cat_ of 4.55×10^3^ s^-1^ (Fig. S3), demonstrating a high affinity to sucrose.

To gain deeper insight into the structural and functional details, the cryo-EM structure of UmSuc2^ΔSP^ was determined at 2.2 Å resolution (Fig. 2A,B; Figs. S4-6). The structure revealed a canonical GH32 architecture consisting of an N-terminal five-bladed catalytic β-propeller and a C-terminal β-sandwich domain. The enzyme assembles as a homodimer (Fig. 2C) and this quaternary arrangement is integral for the proper formation of the active site (Fig. S6). In each protomer, the catalytic pocket is located at the center of the β-propeller but is partially enclosed by structural elements of the adjacent subunit’s β-sandwich, forming a composite cavity (Fig. S4B). This arrangement between both monomers supports that the dimerization is structurally coupled to the formation and stabilization of the catalytic pocket. Superposition of the two monomers demonstrates near-perfect conformational equivalence, with an RMSD of 0.194 Å (Fig. S5D), indicating that both chains adopt indistinguishable global folds and active-site architectures.

**Figure 2:**
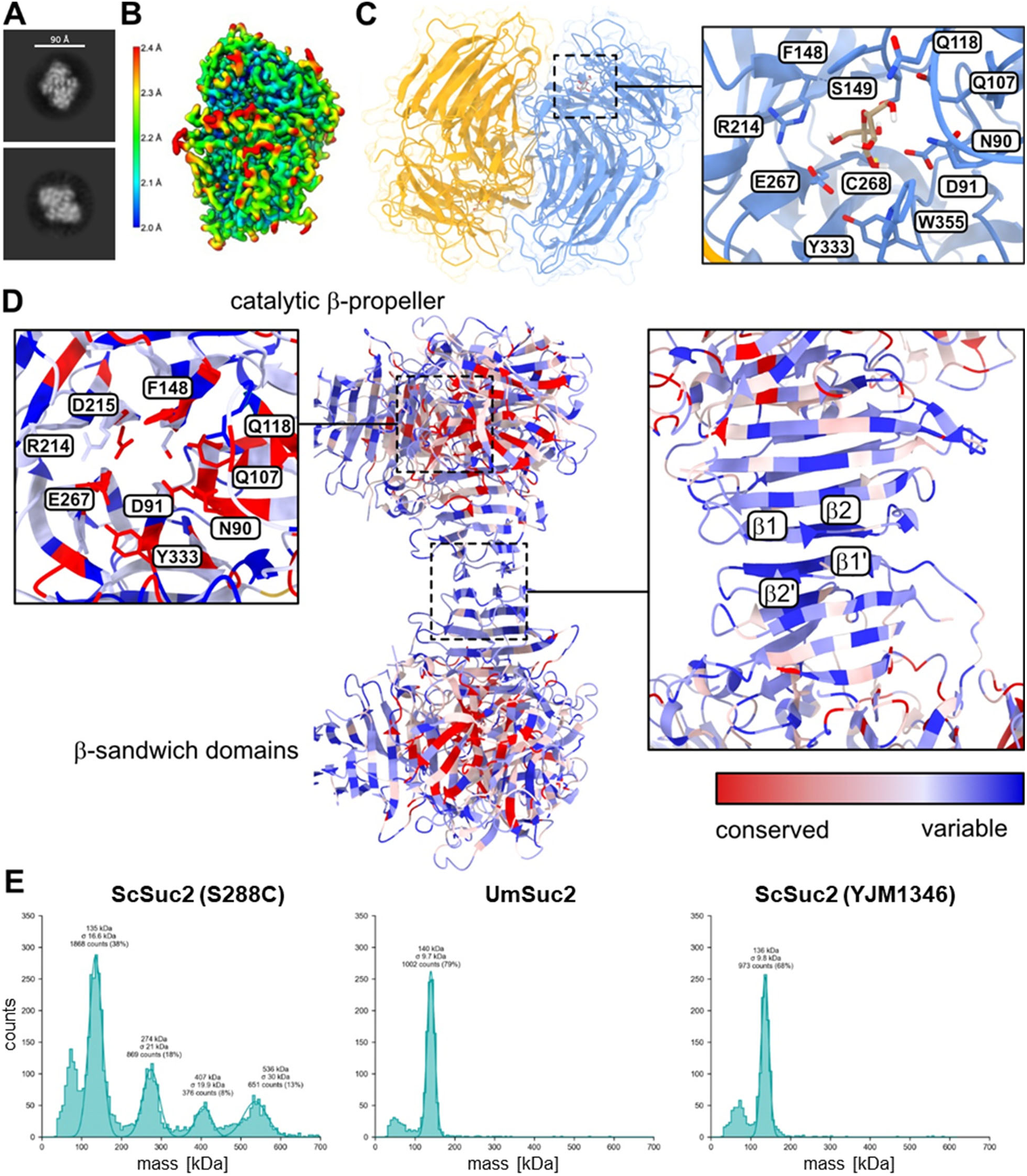
Dimeric structure of recombinant Suc2^ΔSP^. **A,** Representative negative stain images. Scale bar, 90Å. **B**, Local resolution map of the dimer, revealing higher resolution in the well-ordered central core and slightly lower resolution in peripheral surface regions, consistent with increased conformational flexibility at the molecular periphery. **C**, Zoomed-in view of one active site, showing the fructose moiety and surrounding substrate binding residues, illustrating the detailed coordination environment at the catalytic pocket. **D**, Chimera X representation of conservation at dimer:dimer interaction sites of Suc2 variants. AlphaFold structural models of ScSuc2, SoSuc2 and UmSuc2 were used. **E**, Mass photometry reveals higher oligomeric structures up to octamers only for published *S. cerevisiae* S288C Suc2, but not for the *S. cerevisiae* YJM1356 ortholog or UmSuc2.

The atomic model revealed a well-defined density corresponding to fructose in the active site, indicating that the structure captures a product-bound or late catalytic intermediate state. The substrate-binding pockets are formed by thirteen residues lying within 3.6 Å of fructose (Asn90, Asp91, Gln107, Gln118, Phe148, Ser149, Arg214, Glu267, Cys268, Y333 and Trp355), creating a tightly organized interaction network (Fig. 2C). The aromatic residues Phe148, Y333 and Trp355 provide π-stacking and hydrophobic complementarity to the fructosyl ring, while polar and charged residues establish an extensive network of hydrogen bonds and electrostatic interactions that coordinate the glycosidic oxygen and fructosyl hydroxyl groups for hydrolysis. The closest hydrogen-bonding interactions (≤4 Å) involve Asn90, Asp91, and Ser149 residues, which coordinate the entrance and base of the pockets, to orient the substrate into the correct conformation and stabilize the transition state during cleavage of the glycosidic bond. This interaction pattern resembles the polar contact network observed in other structurally characterized invertases that determine sucrose specificity (13, 22).

In contrast to the octameric or tetrameric assembly reported for ScSuc2 or *Purpureocillium lilacinum* invertase, respectively (23), UmSuc2 did not form higher-order complexes under the tested conditions. Instead, its symmetric dimeric architecture closely resembled the organization of secreted dimeric invertase SoInv from *Schwanniomyces occidentalis* (22). Accordingly, structural superposition of UmSuc2 with SoInv (PDB: 3KF5) yielded a Cα RMSD of 0.83 Å (Fig. S5E), demonstrating a strong conservation of the catalytic core. Together, these data indicate that GH32 invertases share a highly conserved catalytic framework embedded in a dimeric assembly, while tolerating limited, species-specific variation in substrate-binding regions. The conserved catalytic geometry, hydrogen-bonding network, and product orientation underscore the evolutionary robustness of the GH32 mechanism.

Based on these findings, we revisited the functional relevance of the higher-order oligomerization described for ScSuc2 (13). Sequence analysis revealed that the catalytic residues are mostly conserved, while the interdimer interactions mediated by strands β1 and β2 of the β-sandwich domain are strongly variable (Fig. 2D). Even more interestingly, Blast-P multi-sequence alignments of ScSuc2 orthologs indicate that these residues are not conserved across different *S. cerevisiae* strains suggesting that higher oligomerization beyond the dimer might not be a general feature but rather an exception in fungal invertases. To test this experimentally, we expressed and purified two ScSuc2 variants: the previously characterized oligomeric form of S288C and a variant from *S. cerevisiae* YJM1356 carrying substitutions at the predicted dimer–dimer interface (Fig. S7). Mass photometry confirmed that UmSuc2 predominantly exists as a dimer, in agreement with our cryo-EM structure (Fig. 2D; Figs. S4 and S5). In contrast, the reference ScSuc2 formed higher-order assemblies, with dimers as the dominant species and tetramers, hexamers and octamers in an apparent equilibrium under the tested conditions (Fig. 2E). Notably, the YJM1356 variant failed to form higher-order oligomers and instead remained predominantly dimeric.

Collectively, these data support the hypothesis that dimers represent the prevalent structural state of fungal invertases, whereas higher-order oligomerization arises from lineage-specific adaptations affecting the interdimer interface.

### 2.2 Canonical proteins for sucrose uptake and hydrolysis are dispensable for axenic growth

To study the physiological relevance of UmSuc2, a gene deletion mutant was generated in *U. maydis* (AB33 suc2Δ). Unexpectedly, the resulting strain still grew on sucrose (Fig. S8), suggesting the presence of additional sucrolytic enzymes. Hence, a second GH32-domain protein named UmSuc1 (UMAG_03605) was investigated as further candidate invertase with 38.03 % sequence identity to ScSuc2 (24, 25) (Figs. 3A, S8A). Despite their limited amino acid sequence homology (Fig. S9A), all proteins share an overall high structural similarity with an r.m.s.d. of 1.1 and 0.7 Å when comparing the AlphaFold model of UmSuc1 and the cryo-EM structure of UmSuc2, respectively, to the crystal structure of ScSuc2 (Fig. S9B) (13). However, notably, UmSuc1 lacks an N-terminal targeting sequence suggesting intracellular sucrose metabolization (Fig. 3A), likely in conjunction with sucrose import via transporter Srt1 (19). However, sucrose-hydrolysing activity was barely detectable in BY4741 *SUC2Δ* cells producing UmSuc1 (Fig. 3B,C). A closer inspection of the active site revealed that the majority of residues is conserved among ScSuc2, UmSuc1 and UmSuc2, except for an arginine that is replaced by a leucine at position 268 in UmSuc1 (Fig. S9A). This suggests that UmSuc1 might only accept sucrose as a minor substrate.

**Figure 3:**
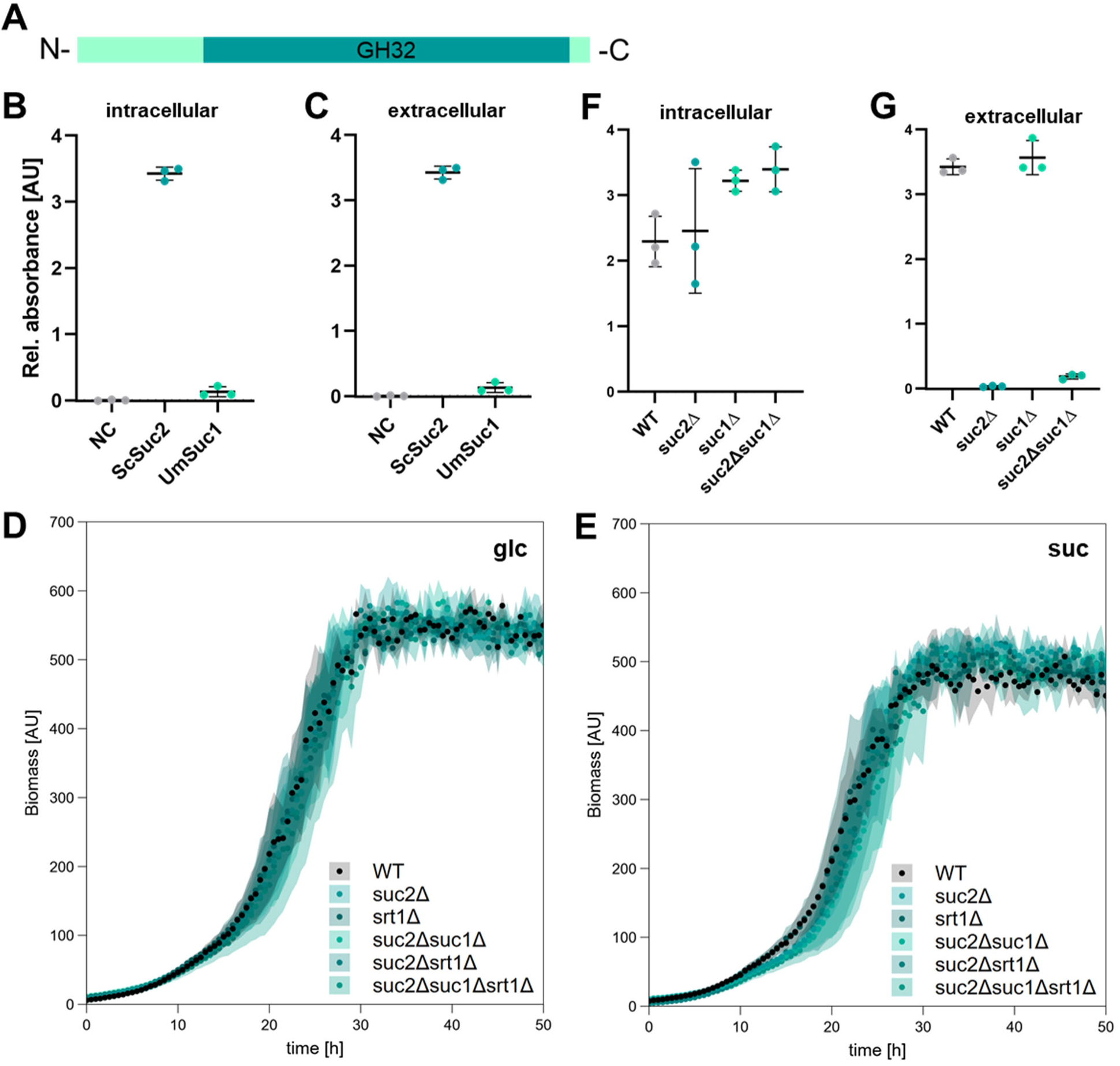
The canonical sucrose metabolism is not essential for axenic growth on sucrose. **A,** Schematic representation of Suc1 domain architecture. **B**, **C**, DNS assays with sucrose as a substrate of tester strain *S. cerevisiae* BY4741 *SUC2Δ* derivatives producing UmSuc1 and ScSuc2 (positive control). All strains were grown in liquid cultures with glucose until the carbon source was depleted. Then, DNS assays were conducted using cell extracts (**B**) or cell-free culture supernatants (**C**) and sucrose as a substrate to detect reducing sugars resulting from hydrolysis of the disaccharide. Detection of reducing sugars is indicative for sucrose hydrolysis. NC, negative control using BY4741 *SUC2Δ* without additional genes. Assays were conducted in biological triplicates. Error bars depict standard deviation. **D**, **E,** Growth of *U. maydis* laboratory strain AB33 (WT) and the indicated deletion mutants on glucose (**D**) and sucrose (**E**). The diagrams show backscatter data of indicated strains obtained in a micro-scale cultivation visualizing cell propagation. The assay was conducted in biological triplicates. Error bars depict standard deviation. AU, artificial units. Growth rates of averaged data are shown below. **F,G,** DNS assays conducted with cell extracts (**F**) or culture supernatants (**G**) from indicated *U. maydis* AB33 derivatives pre-grown on sucrose as single carbon source.

To analyse the role of these proteins for sucrose metabolization in *U. maydis*, we generated single deletion mutants of *suc1*, *suc2* and *srt1* as well as double and triple mutants in all possible combinations. However, online growth monitoring revealed that all strains grew comparable on glucose and sucrose, with similar average growth rates between 0.14 h^-1^ and 0.17 h^-1^ and similar final biomass (Fig. 3D, E; Table S1). While sucrose-hydrolysing activity was diminished in the culture supernatant of strains lacking Suc2 (Fig. 3F), the disaccharide was still hydrolyzed in cell extracts lacking Suc2 or even Suc1 and Suc2 (Fig. 3G), indicating the presence of further intracellular enzymes for sucrose hydrolysis.

Together, we established UmSuc2 as a functional invertase with extracellular activity, whereas intracellular Suc1 displayed only marginal sucrose-hydrolyzing capacity. Strikingly, however, neither deletion of both GH32 enzymes nor simultaneous removal of the transporter Srt1 impaired growth on sucrose in the yeast form, demonstrating that canonical invertase-dependent sucrose metabolism is dispensable during axenic growth. These findings indicate that *U. maydis* can compensate for the loss of the canonical proteins of the sucrose metabolism and relies on an alternative sucrose utilization mechanism in this growth stage.

### 2.3 Repurposed enzymes from maltose metabolism are employed for axenic growth on sucrose

Based on the assumption that further sucrose-hydrolyzing enzymes and transporters must be present in *U. maydis*, the search for potential candidates was expanded. In *S. cerevisiae*, isomaltase Ima1 and maltase Mal12 also accept sucrose as a substrate (5, 26). Thus, we hypothesized that a similar scenario could exist in *U. maydis*. Indeed, it possesses two homologs with predicted GH13 domains, maltase UmMal12 (UMAG_03692) and isomaltase UmIma1 (UMAG_15026). The *U. maydis* homologs share high sequence identity of 44 % and 45 % with ScIma1 and ScMal12, respectively (Fig. S10A). Accordingly, structure predictions also reveal a high conservation with respect to the *S. cerevisiae* homologs (Fig. S10B). Besides the two enzymes, a homolog of *S. cerevisiae* Agt1, a maltose transporter potentially accepting sucrose as a substrate (termed UmAgt1; UMAG_05972) (27) is present.

To verify if UmIma1 and UmMal12 can hydrolyse sucrose, we again employed *S. cerevisiae* tester strain BY4741 *SUC2Δ* (Fig. 4A). Indeed, sucrose hydrolysis were detected in cell extracts when UmIma1 was present, suggesting that this enzyme is capable of accepting sucrose as a substrate. By contrast, Mal12 producing strains only revealed low sucrolytic activity, suggesting a rather weak turnover of sucrose. In essence, these data demonstrate that *U. maydis* orthologs of proteins known from *S. cerevisiae* maltose metabolism also efficiently use sucrose as substrates.

**Figure 4:**
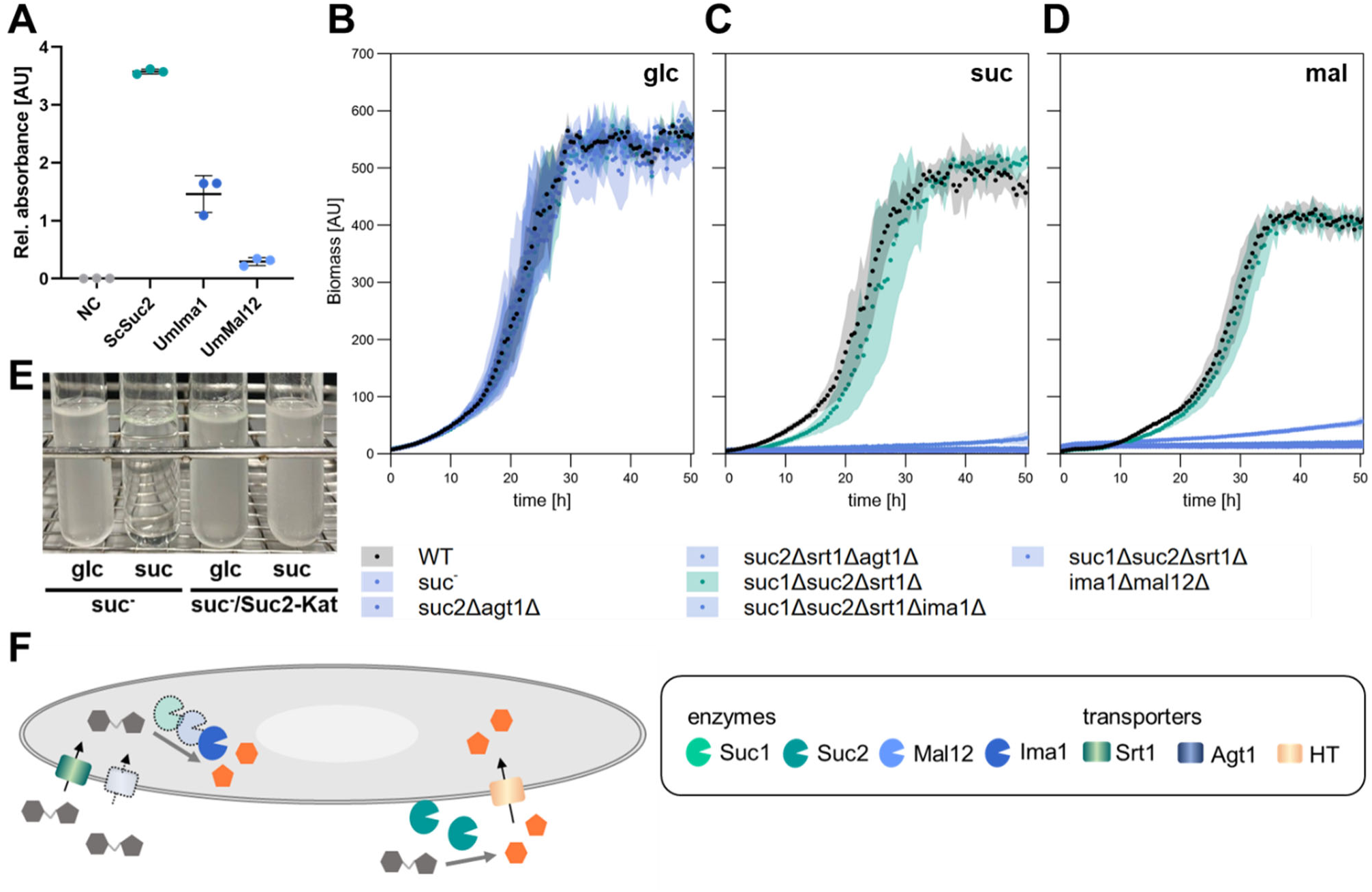
Sucrose utilization in axenic growth relies on repurposed maltose metabolism machinery. **A,** DNS assays with sucrose as a substrate using cell extracts of reporter strain *S. cerevisiae* BY4741 *SUC2Δ* derivatives producing ScSuc2, UmIma1 and UmMal12. Detection of reducing sugars is indicative for invertase activity. Assays were conducted in biological triplicates. Error bars depict standard deviation. **B-D,** Growth of *U. maydis* laboratory strain AB33 (WT) and the indicated deletion mutants on glucose (**B**), sucrose (**C**) and maltose (**D**). The diagrams show backscatter data of indicated strains obtained in a micro-scale cultivation visualizing cell propagation. The assay was conducted in biological triplicates. Error bars depict standard deviation. AU, artificial units. Glc, glucose; suc, sucrose; mal, maltose. **E,** Conditional growth by public metabolism in AB33 suc^-^ (lacking all genes needed for sucrose metabolization), complemented with *suc2-kat* controlled by the native promoter. **F,** Model of sucrose metabolism during axenic growth of yeast-like cells. HT, hexose transporter. Orange, hexoses; grey, sucrose. For enzyme and transporter specifications, see legend.

To test the functional relevance of UmMal12 and UmIma1 in the homologous system, a set of *U. maydis* AB33 derivatives lacking different combinations of these enzymes, as well as a quadruple deletion strain were generated. Growth on glucose did not show any significant differences between the different strains with similar growth rates and final biomass (Fig. 4B, S8; Table S2). By contrast, sucrose as single carbon source resulted in a severe growth retardation in a strain carrying deletions of *suc1*, *suc2* and *ima1*, and growth was completely ablated in the quadruple deletion mutant with additional deletion of *mal12* (Fig. 4C; Table S2). This confirms that Ima1 strongly contributes to sucrose acquisition in the yeast form. The role of Agt1 was addressed using a *srt1*/*agt1*/*suc2* triple deletion strain lacking both sucrose transporter candidates and the only secreted invertase to avoid extracellular hydrolysis with subsequent uptake of hexoses. This strain also failed to grow on sucrose, suggesting that we identified the complete set of sucrose transport proteins employed for axenic growth (Fig. 4C; Table S2). As the proteins were partially adapted from maltose metabolism, growth was also characterized on maltose (Fig. 4D; Table S2). Indeed, strains lacking *ima1* and *mal12* or *agt1* also did not grow on maltose, suggesting that the ability of hydrolyzing and transporting this disaccharide has been retained.

The preceding analyses indicates that both extra- and intracellular sucrose metabolism are employed by *U. maydis*, and the intracellular metabolism can fully compensate for the absence of secreted Suc2 under the tested laboratory conditions. To test if this is also true for the opposite scenario, we generated AB33 suc^-^ lacking all six proteins for sucrose import and hydrolysis and complemented it with a *suc2:mKate2* fusion controlled by the native *suc2* promoter. Both strains grew identical on glucose but only the complemented strain AB33 suc^-^/Suc2-Kat grew on sucrose, demonstrating its ability to purely rely on extracellular hydrolysis for sucrose acquisition (Fig. 4E).

Collectively, we uncovered the full set of enzymes required for axenic yeast-like growth on sucrose (Fig. 4F). Our data support the idea of a dual strategy, enabling the fungus to employ canonical extracellular invertase UmSuc2 or import sucrose for hydrolysis with repurposed intracellular enzymes derived from maltose metabolism. While Ima1 and Agt1 are the dominant determinants of intracellular metabolism, Mal12 and Suc1 do not contribute significantly under the tested conditions (Fig. 4F).

### 2.4 Sucrose import is key to efficient plant infection

In line with our findings, published transcriptional profiling data (28) indicate that *ima1* and *agt1* are highly expressed in axenic culture while *mal12* transcripts are barely detectable. By contrast, *suc2* and *srt1* transcripts showed low abundance in axenic culture, but specific induction during plant infection (Fig. S11). *suc1* transcripts showed an overall low abundance with slightly higher values during axenic growth. These transcriptional profiles are consistent with a context-dependent deployment of sucrose acquisition modules, with *ima1* and *agt1* used for axenic growth on sucrose and *suc2* and *srt1* employed during plant infection.

To verify the relevance of these sucrose acquisition modules during plant infection, corresponding deletion mutants were generated in the solopathogenic *U. maydis* strain SG200 (Fig. 5). This haploid strain infects maize independently of mating and serves as a robust model for virulence analyses (15). Consistent with previous reports, deletion of *srt1* resulted in reduced pathogenicity (19), confirming the importance of high-affinity sucrose uptake during infection. In contrast, elimination of *agt1* did not significantly affect symptom development, indicating that the major axenic sucrose transporter is dispensable *in planta*. Strikingly, the *srt1/agt1* double mutant displayed strongly attenuated virulence, demonstrating that sucrose uptake for intracellular metabolization is critical for efficient infection. Additional deletion of *suc2*, caused only a minor further reduction in symptom development. These findings indicate intracellular sucrose acquisition via Srt1 constitutes the dominant strategy for carbon uptake *in planta*. The residual pathogenicity observed in transporter-deficient strains suggests that *U. maydis* can access alternative carbon sources like glucose or malate (29) within the host, albeit with strongly reduced efficiency (Fig. 5).

**Figure 5:**
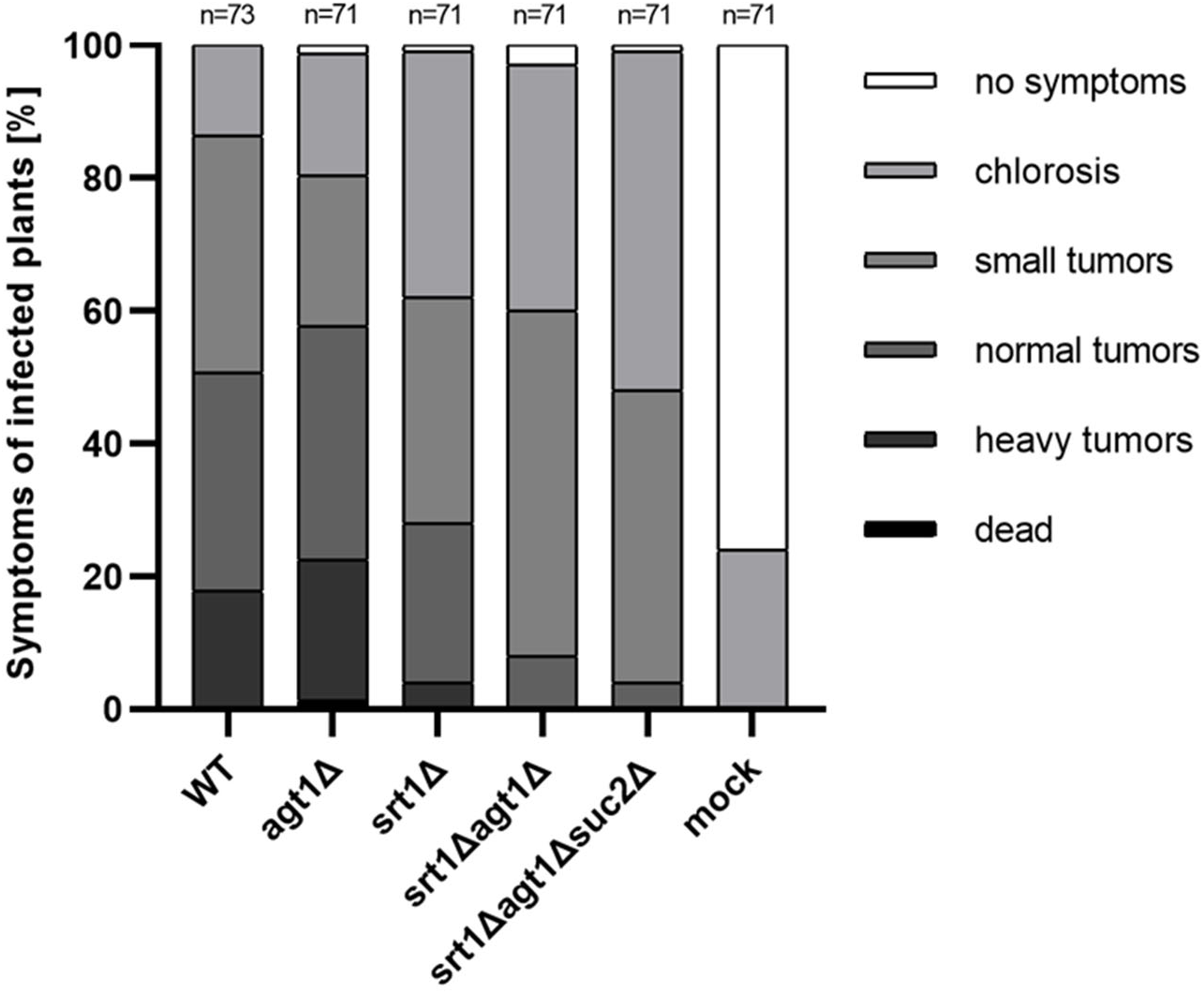
Sucrose metabolism is critical for maize infection. Infection assays with SG200 and the indicated derivatives to characterize the ability to infect maize plants in the absence of sucrose transporters, private sucrose metabolism and sucrose metabolism. Darker colors reflect stronger symptoms while lighter colors represent mild symptoms (see legend). Ranking was conducted 12 dpi. n= number of infected plants. Infection assays were conducted in biological triplicates.

Together, these results reveal a clear functional separation between saprotrophic and pathogenic sucrose utilization strategies. Whereas axenic growth relies on secreted Suc2 and a repurposed maltose module, efficient colonization of maize requires high-affinity sucrose uptake mediated by Srt1.

## 3 Discussion

In this study, we uncover the complete enzyme repertoire for sucrose metabolization in the biotrophic fungal pathogen *U. maydis*, characterized by an unexpected flexibility, re-purposed enzymes and context-dependent employment of differential protein subsets. This metabolic flexibility enables the fungus to adapt to its different niches, yeast-like growth and biotrophic infection. While *U. maydis* is generally capable of both, extracellular and intracellular sucrose utilization, the sucrose metabolism in the infectious stage is clearly streamlined for efficient sucrose uptake. Our observations in *U. maydis* are contrasting findings in *S. cerevisiae* that employs mainly extracellular sucrose metabolism mediated by invertase ScSuc2 in sucrose-rich environments (2). This difference might be attributed to the different lifestyles as a ubiquitously found nomadic microbe (30) versus a specialized plant pathogen with a very narrow host range (31).

### 3.1 Canonical secreted invertase constitutes a dimer partially retained at the cell boundary

*U. maydis* contains a canonical secreted invertase, UmSuc2, for extracellular sucrose hydrolysis. In yeast-like growing cells, we provide evidence for a dual localization: Invertase activity attributed to UmSuc2 is detected in the cell-free culture supernatant. However, based on fluorescence microscopy the enzyme is also partially retained at the mother cell boundary. Daughter cells lack this peripheric signal, suggesting that the secreted protein passes through their cell walls, probably due to the differing cell wall composition in expanding buds. By contrast, a Suc2-Gfp fusion protein localized to both mother and daughter cells in *S. cerevisiae* (32). Of note, the cell-bound signal for UmSuc2 disappears in protoplasts of *U. maydis*, suggesting that the cell wall constitutes a critical barrier for its release. Similar observations were made in biochemical fractionation experiments in the filamentous ascomycete *Aspergillus nidulans* (33).

Our biochemical data established that UmSuc2 is an acidic invertase with a K_m_ of 53 mM, a V_max_ of 2.4 mM/min, and a turnover K_cat_ of 4.55×10^3^ s^-1^. The relatively low K_m_ value indicates high affinity for sucrose and is comparable to those reported for fungi with sequence similarities, e.g., *Penicillium aurantiogriseum* (65.28 mM), *Uromyces fabae* (27.1 mM), and *Aspergillus niger* (15.82 mM) (34, 35). Interestingly, recombinant UmSuc2 constitutes as a dimer, similar to *S. occidentalis* invertase SoInv (22), and both proteins share a high degree of structural similarity (Fig. 2 and Fig. S5). By contrast, *S. cerevisiae* S288C Suc2 assembles into higher-oligomeric structures, and octamers are thought to represent the major extracellular form (13). The limited conservation of residues mediating dimer:dimer interactions, together with our observation that recombinant ScSuc2 from the clinical isolate YJM1356 (36) predominantly forms dimers, suggests that higher-order oligomerization evolved only in distinct lineages rather than representing a general property of fungal invertases. This raises the possibility that formation of higher-order oligomers may be favored only under specific physiological or ecological conditions, such as repeated growth in sucrose-rich environments. Notably, the octameric structure in conjunction with extensive N-glycosylation has also been hypothesized to prohibit the passage through the yeast cell wall through structural constraints (12, 13)(18,19). In *U. maydis*, the partial retainment of the dimeric Suc2-Gfp fusion protein, a protein complex of about 190 kDa, indicates that octamerization is probably not required for trapping the enzyme at the cell boundary.

Interestingly, the assembly into dimers versus higher oligomers in SoSuc2 and ScSuc2, respectively, influences the active site architecture. This results in differential substrate specificities, with ScSuc2 preferring sucrose as major substrate, whereas SoSuc2 also efficiently hydrolyzes fructans like nystose (22). Hence, UmSuc2 likely provides a similar substrate flexibility, not only efficiently hydrolyzing sucrose but also fructans. This could provide advantages in the race for carbon during axenic growth, i.e., in the context of the soil or root microbiome. Maize roots for example possess fructan exohydrolase (37), suggesting the presence of fructose oligomers in the rhizosphere.

Based on the finding that octameric invertase has to our knowledge only been found in S288C until now, but is likely not present in neither related Ascomycete or basidiomycete yeasts (38), nor in *U. maydis*, one may speculate that formation of higher-order oligomers and hence the preference for sucrose over fructans in distinct lineages of *S. cerevisiae* ScSuc2 might be an evolutionary adaptation to sucrose-rich environments due to the long history of co-existence with humans. While such “domesticated” yeast strains are employed in human-shaped environments during food and beverage fermentation for ages, wild isolates show a wide distribution of habitats, ranging from plant exudates, flower nectars and rotten wood to fruits and bark (2, 39), where they may rely on a higher metabolic flexibility.

### 3.2 *U. maydis* sucrose metabolism reveals unexpected flexibility

*U. maydis* follows a dual strategy with secreted Suc2 for extracellular as well as Ima1 and Agt1 for intracellular hydrolysis during yeast-like growth on sucrose. Two enzymes invented for maltose metabolization hence play a key role during axenic growth, where they equally accept maltose and sucrose as substrates (Fig. 6). Notably, growth on sucrose does not need maltose as inductor, as observed for *S. cerevisiae* (26). Little is known about the natural context of yeast-like growth in *U. maydis*. Teliospores frequently develop in decaying corn cobs, where starch released from kernels is likely a major carbon source. Albeit potential enzymes are present, degradation of starch has not been observed in *U. maydis* (24, 40). Still, the resulting monosaccharide maltose may represent an abundant environmental nutrient during yeast-like growth, potentially explaining the importance of the maltose utilization machinery under axenic conditions. Sucrose, the main transport sugar in living plants, is thought to be rapidly depleted in decaying plant tissue by competing microbes. We therefore speculate that *U. maydis* is prepared for efficient uptake of both sucrose and maltose, thereby increasing metabolic flexibility and competitiveness during growth in microbe-rich decaying plant material.

**Figure 6:**
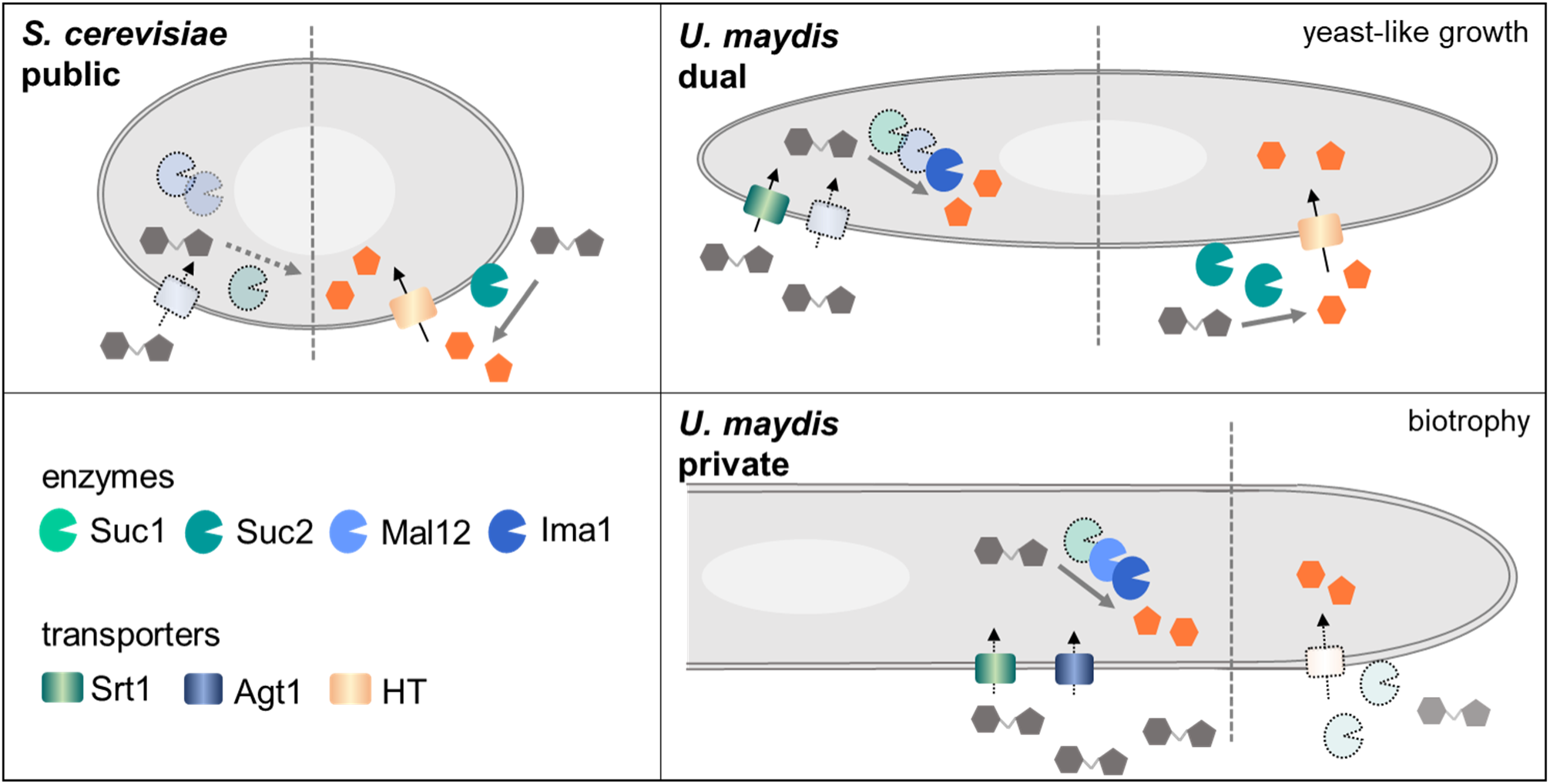
Comparative model of sucrose metabolism in *S. cerevisiae* and *U. maydis*. *S. cerevisiae* mainly employs secreted invertase ScSuc2 for extracellular sucrose hydrolysis as canonical enzyme. Resulting monosaccharides constitute public goods. By contrast, *U. maydis* uses a broader repertoire of enzymes. During yeast-like growth it relies on a dual strategy, with external hydrolysis by public invertase UmSuc2 or via private metabolism through sucrose uptake and intracellular hydrolysis. Proteins dedicated to maltose metabolism can alternatively cope for acquiring sucrose in both organisms, however, this pathway has evolved as a major route in yeast-like cells of *U. maydis*, while it is only supporting slow growth of *S. cerevisiae* in the absence of the canonical enzyme. Transparent enzymes or transporters depict proteins that are not essential for growth on sucrose at standard conditions. Dashed lines separate private (left) and public metabolism (right).

*In planta*, *U. maydis* employs a combination of the high-affinity sucrose transporter Srt1 and repurposed Agt1 for efficient carbon acquisition. Hence, cells privatize their sucrose metabolism during biotrophy, importing sucrose via membrane transporters and subsequent intracellular hydrolysis for fueling glycolysis with the resulting monosaccharides (Fig. 6). This differs from the preference of public sucrose metabolism described for *S. cerevisiae*. Here, maltose transporters and intracellular enzymes from maltose metabolism in principle accept sucrose as substrates. However, the expression of the enzymes is not induced by sucrose, hence their abundance is usually too low for efficient metabolization in the absence of extracellular ScSuc2 (26). We hypothesize that these differences reflect niche-related adaptations of the genetic repertoire during evolution of this intimate plant-pathogen interaction. Privatization during plant infection of *U. maydis* might be a result of arms race where the fungus uses the high-affinity sucrose transporter Srt1 together with Agt1 to quickly acquaint the disaccharide for efficient *in planta*-growth. In line with that, maize cell-wall invertase is induced during infection (28), hence it would be advantageous for the fungus to “steal” available sucrose for internal metabolization. At the same time, this contributes to the artificial source-sink transition, by inducing an increased flow of sucrose into infected tissue (16, 17, 41).

In conclusion, our study reveals an astonishing metabolic flexibility for sucrose acquisition, that is likely adapted to the biphasic lifestyle of the fungal plant pathogen. Our observations indicate that *U. maydis* evolved a complex and powerful strategy, likely allowing for conditional switching between public and private sucrose metabolism in the yeast stage. While invertase secretion might be beneficial for scouting at nutrient deprivation, privatization of sucrose hydrolysis is important for evasion of the plant immune system and efficient sucrose acquisition *in planta*. Moreover, this metabolic flexibility might also yield benefits in the race for carbon during competition with other (micro)organisms, both in the rhizosphere and *in planta*.

## 4 Materials and methods

### 4.1 Accession numbers

Proteins that are studied in this work are listed in uniprot database (https://www.uniprot.org; accessed 11/05/2025) with the following accession numbers: *S. cerevisiae* S288C Suc2 (ScSuc2): P00724; *U. maydis* Suc1 (umag_03605; UmSuc1): A0A0D1C4E6; *U. maydis* Suc2 (umag_01945; UmSuc2): A0A0D1E3W7; *U. maydis* Srt1 (umag_02374; UmSrt1): A0A0D1E1N5; *U. maydis* Agt1 (umag_05972; UmAgt1): A0A0D1BW32; *U. maydis* Ima1 (umag_15026; UmIma1): A0A0D1CN93; *U. maydis* Mal12 (umag_03692; UmMal12): A0A0D1DUG9.

### 4.2 Molecular biology methods

All plasmids (pUMa/pUX vectors) generated in this study are listed in Table S3. All plasmids were obtained using standard molecular biology methods established for *U. maydis* including restriction ligation, Golden Gate and Gibson cloning (42, 43). Genomic DNA of *U. maydis* strain UM521 was used as template for PCR reactions. The genomic sequence for this strain is stored at the EnsemblFungi database. Restriction enzymes for cloning were purchased from NEB (Ipswich, MA, USA).

### 4.3 Strain generation

All *U. maydis* strains were generated by homologous recombination yielding genetically stable strains based on the laboratory strain AB33 or solopathogenic strain SG200 (15, 44) (Table 1). For all genetic manipulations, protoplasts were generated and transformed with linear DNA fragments (or free-replicating plasmids) using established protocols (45). All strains were verified by Southern blot analysis (deletion/complementation strains) or diagnostic PCR (FRT recycling). For *in locus* modifications the up- and downstream flanks were amplified as probes. When other loci were used for insertion (46), standard deletion flanks were employed (Table 2). Flippase (FLP)-mediated resistance marker recycling was conducted according to published protocols (47) yielding marker-free strains. Different FRT sites were used to avoid recombination between remaining FRT scars.

**Table 1:** *U. maydis* strains used and generated in this study. ^R^, resistance markers were removed by Flp-mediated recombination (47).

| ***U. maydis* strain** | **Internal strain collection number<sup>1</sup>** | **Parental strain** | **Vector<sup>2</sup>** | **Genotype** | **Reference** |
| --- | --- | --- | --- | --- | --- |
| AB33 | UMa133 | FB2 | - | *a2 Pnar: bW2bE1 PhleoR* | (44) |
| AB33 *suc2Δ* | UX0029 | AB33 | pUX54 | *a2 Pnar: bW2bE1 PhleoR umag_01945Δ_HygR* | This work. |
| AB33 *suc2Δ* | UX0030 | UX0029 | pUMA1446 | *a2 Pnar: bW2bE1 PhleoR umag_01945Δ* | This work. |
| AB33 *srt1Δ* | UX0041 | AB33 | pUX52 | *a2 Pnar: bW2bE1 PhleoR umag_02374Δ_HygR* | This work. |
| AB33 *suc2Δ*<br>*suc1Δ* | UX0049 | UX0030 | pUX53 | *a2 Pnar: bW2bE1*<br>*PhleoR umag_01945Δ*<br>*umag_03605Δ_HygR* | This work. |
| AB33 *suc2Δ*<br>*suc1Δ* | UX0050 | UX0049 | pUMa1446 | *a2 Pnar: bW2bE1*<br>*PhleoR umag_01945Δ*<br>*umag_03605Δ* | This work. |
| AB33 *suc2Δ*<br>*suc1Δ srt1Δ* | UX0051 | UX0050 | pUX52 | *a2 Pnar: bW2bE1*<br>*PhleoR umag_01945Δ*<br>*umag_03605Δ*<br>*umag_02374Δ_HygR* | This work. |
| AB33 *suc2Δ*<br>*srt1Δ* | UX0090 | UX0030 | pUX52 | *a2 Pnar: bW2bE1*<br>*PhleoR umag_01945Δ*<br>*umag_02374Δ_HygR* | This work. |
| AB33 *suc2Δ*<br>*suc1Δ ima1Δ* | UX0122 | UX050 | pUX149 | *a2 Pnar: bW2bE1*<br>*PhleoR umag_01945Δ*<br>*umag_03605Δ*<br>*umag_15026Δ_HygR* | This work. |
| AB33 *suc2Δ*<br>*suc1Δ ima1Δ*<br>*mal12Δ* | UX0123 | UX0122 | pUX181 | *a2 Pnar: bW2bE1*<br>*PhleoR umag_01945Δ*<br>*umag_03605Δ*<br>*umag_15026Δ_HygR*<br>*umag_03692Δ_NatR*<br>Precursor for UX0124 | This work. |
| AB33 *suc2Δ*<br>*suc1Δ ima1Δ*<br>*mal12Δ<sup>R</sup>* | UX0124 | UX0123 | pUMa1446 | *a2 Pnar: bW2bE1*<br>*PhleoR umag_01945Δ*<br>*umag_03605Δ*<br>*umag_15026Δ*<br>*umag_03692Δ*<br>Resistance markers<br>HygR/NatR removed. | This work. |
| AB33 *suc2Δ*<br>*srt1Δ agt1Δ* | UX0125 | UX0090 | pUX173 | *a2 Pnar: bW2bE1*<br>*PhleoR umag_01945Δ*<br>*umag_02374Δ_HygR*<br>*umag_05972Δ_NatR* | This work. |
| AB33 *suc2Δ*<br>*suc1Δ ima1Δ*<br>*mal12Δ srt1Δ* | UX0126 | UX0124 | pUX52 | *a2 Pnar: bW2bE1*<br>*PhleoR umag_01945Δ*<br>*umag_03605Δ*<br>*umag_15026Δ* *umag_03692Δ*<br>*umag_02374Δ_HygR* | This work. |
| AB33 *suc2Δ*<br>*agt1Δ* | UX0155 | UX0030 | pUX173 | *a2 Pnar: bW2bE1*<br>*PhleoR*<br>*umag_01945Δ_HygR*<br>*umag_05972Δ_NatR* | This work. |
| AB33 *suc<sup>-</sup>*<br>(AB33 *suc2Δ*<br>*suc1Δ ima1Δ*<br>*mal12Δ srt1Δ*<br>*agt1Δ*) | UX0156 | UX0126 | pUX173 | *a2 Pnar: bW2bE1*<br>*PhleoR umag_01945Δ*<br>*umag_03605Δ*<br>*umag_15026Δ*<br>*umag_03692Δ*<br>*umag_02374Δ_HygR*<br>*umag_05972Δ_NatR*<br>Precursor for UX0157. | This work. |
| AB33 *suc<sup>-R</sup>*<br>(AB33 *suc2Δ*<br>*suc1Δ ima1Δ*<br>*mal12Δ srt1Δ*<br>*agt1Δ<sup>R</sup>*) | UX0157 | UX0156 | pUMa1446 | *a2 Pnar: bW2bE1*<br>*PhleoR umag_01945Δ*<br>*umag_03605Δ*<br>*umag_15026Δ*<br>*umag_03692Δ*<br>*umag_02374Δ*<br>*umag_05972Δ*<br>Resistance markers<br>HygR/NatR removed. | This work. |
| AB33<br>*suc2Δ/Suc2-*<br>Kat | UX0320 | AB33 | pUX0487 | *a2 Pnar: bW2bE1*<br>*PhleoR umag_01945Δ*<br>*Psuc2:suc2-*<br>*mKate2_HygR* | This work. |
| AB33 *suc<sup>-</sup>*<br>/Suc2-Kat<br>(AB33 *suc2Δ*<br>*suc1Δ ima1Δ*<br>*mal12Δ srt1Δ*<br>*agt1Δ/Suc2-*<br>Kat) | UX0450 | UX0157 | pUX0487 | *a2 Pnar: bW2bE1*<br>*PhleoR umag_01945Δ*<br>*umag_03605Δ*<br>*umag_15026Δ*<br>*umag_03692Δ*<br>*umag_02374Δ_HygR*<br>*umag_05972Δ_NatR*<br>*Psuc2:suc2-*<br>*mKate2_HygR* | This work. |
| SG200 | UMa3409 | FB2 |  | *a1mfa2bE1bW2* | (15) |
| SG200 *agt1Δ* | UX0223 | SG200 | pUX476 | *a1mfa2bE1bW2*<br>*umag_05972_NatR* | This work. |
| SG200 srt1Δ | UX0224 | SG200 | pUX52 | *a1mfa2bE1bW2*<br>*umag_02374_HygR* | This work. |
| SG200 agt1Δ<br>srt1Δ | UX0225 | UX0224 | pUX476 | *a1mfa2bE1bW2*<br>*umag_02374_HygR*<br>*umag_05972_NatR* | This work. |
| SG200 agt1Δ<br>srt1Δ | UX0226 | UX0225 | pUMa1446 | *a1mfa2bE1bW2*<br>*umag_02374*<br>*umag_05972* | This work. |
| SG200 agt1Δ<br>srt1Δ suc2Δ | UX0227 | UX0226 | pUX54 | *a1mfa2bE1bW2*<br>*umag_02374*<br>*umag_05972*<br>*umag_01945_HygR* | This work. |
| SG200<br>suc2Δ/ Suc2-Kat | UX0321 | SG200 | pUX487 | *a1mfa2bE1bW2*<br>*umag_01945Δ*<br>*Psuc2:suc2-*<br>*mKate2_HygR* | This work. |

<sup>1</sup> Internal strain collection numbers (UMa/Ux codes)
<sup>2</sup> Plasmids generated in our Institute are stored in a plasmid collection and termed pUMa or pUx plus a 4-digit number as identifier.

**Table 2:**
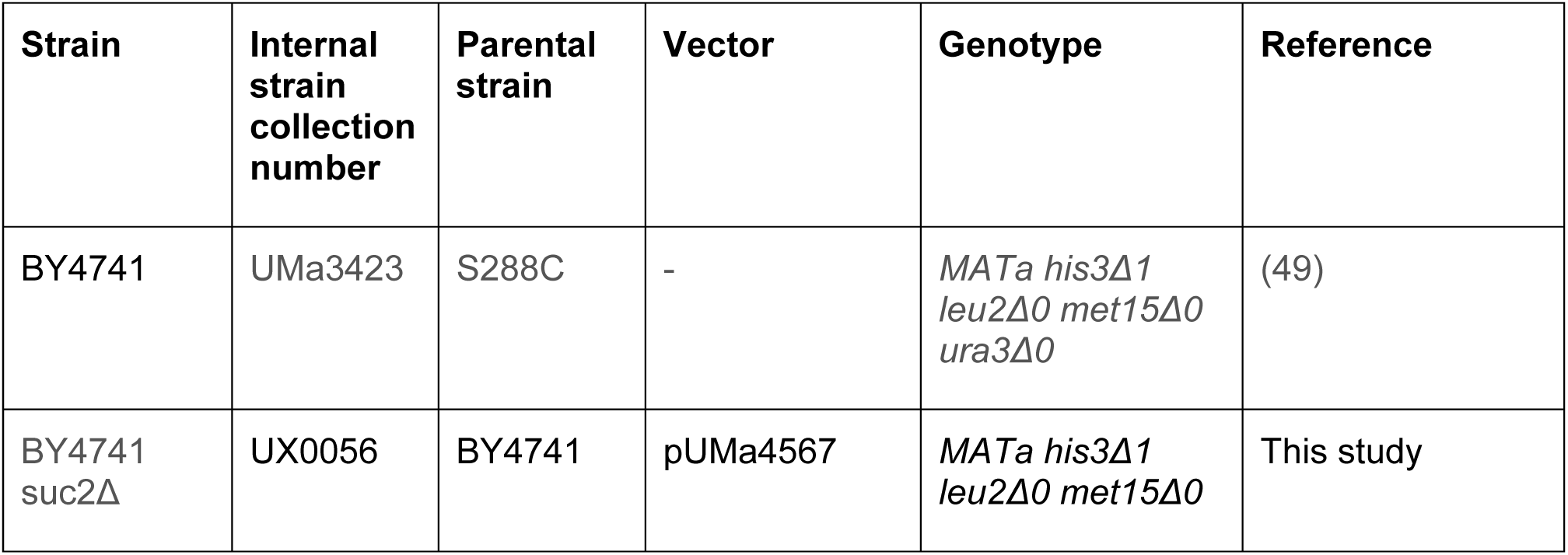
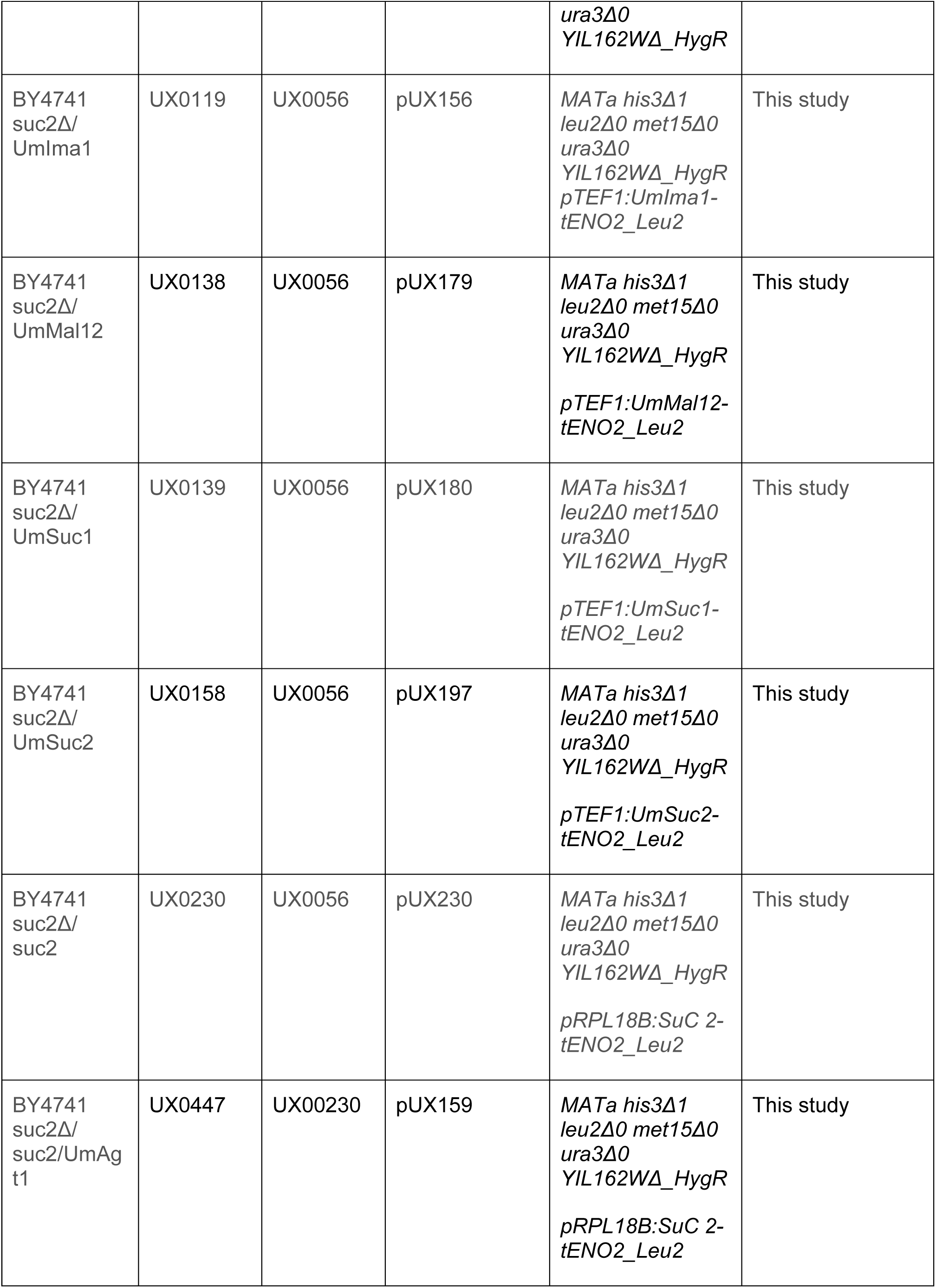
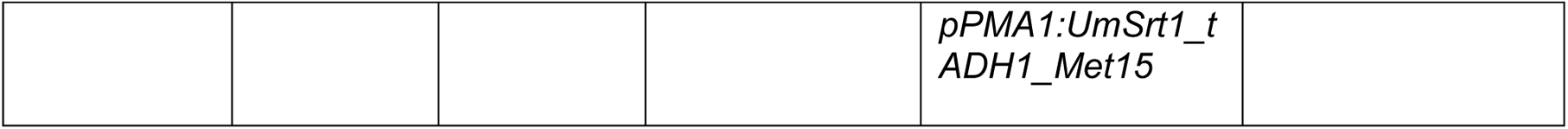
*S. cerevisiae* strains used and generated in this study.

*S. cerevisiae* strains were generated by complementing auxotrophies (*his3Δ1 leu2Δ0 met15Δ0 ura3Δ0)* present in the progenitor strain BY4741, an S288C derivative. Alternatively, a hygromycin resistance cassette was employed (HygR) for selection. PEG-mediated strain transformation was conducted using competent cells according to published protocols (48). All strains were verified by diagnostic PCR (Table 2).

### 4.4 Cultivation, online monitoring and growth rate determination

*U. maydis* strains were grown at 28 °C in complete medium supplemented with 1% (w/v) glucose (CM-glc) if not described differently by inoculation from a fresh plate. Solid media were supplemented with 2% (w/v) agar agar. For experiments in a BioLector microbioreactor (m2p-labs), a pre-culture was grown in CM-glc. On the next day, cells were adjusted to an optical density of 600 nm (OD_600_) of 0.2 in Verduyn medium supplemented with 1 % glucose (or different saccharide as indicated) and incubated for 6 h on a rotary shaker at 28°C. Cells were then washed once with sterile H_2_O_bidest_ and resuspended in Verduyn with the respective carbons source (30 mM sucrose, 30 mM maltose, 60 mM glucose). MTP-48-B flower plates were inoculated with 1.5 ml cell of these cell suspensions per well (in technical duplicates) and incubated at 28°C at 1,000 rpm. Backscattered light was measured with a 620 hm filter to determine biomass. For data analysis, a blank (pure medium) was averaged and subtracted from the other wells with the similar medium. All experiments were conducted in three biological replicates. All experiments were conducted in three biological replicates. Exponential-phase specific growth rates were calculated from averaged scattered-light biomass measurements (mean of three replicates) over the 5–25 h interval. The analysis, including all equations, is documented in a Jupyter notebook (Supplementary file 1; https://github.com/Tanvir96rwth/Growth_rate_analysis.git).

### 4.5 Determination of reducing sugars

Reducing sugars (glucose, fructose) were determined using 3,5-dinitrosalicylic acid (DNS) assays (50). For *S. cerevisiae* experiments, pre-cultures in 5 ml YPD (10 g/l yeast extract, 20 g/L peptone, 20 g/l D (+)-glucose monohydrate were inoculated with the respective strain and incubated overnight at 28°C on a rotary shaker. The 10 ml main culture was inoculated in YNB (amino acid and nitrogen base) supplemented with 20 g/l glucose using the pre-culture to reach a starting OD_600_ of 0.1, incubated at 28°C and 200 rpm (Throw 25 mm) for 40 hours. For *U. maydis*, pre-cultures in 5ml CM-glucose were inoculated with the respective strain and incubated overnight at 28°C in a rotary shaker. For the main culture 10 ml of Verduyn – glucose (10g/l) or sucrose (10g/l) were inoculated using the pre-culture to reach a starting OD_600_ of 0.1, incubated at 28°C and 200 rpm (Throw 25 mm) for 48 hours.

All cultures were tested for the absence of remaining free glucose using glucose test stripes for urine analysis (Medi-Test Glucose, Macherey-Nagel). Subsequently, for harvest, the optical density at 600 nm was documented and then, cells were centrifuged for 3 min in 15 ml tubes at 3,409 x g). The supernatant was transferred to a new tube and stored on ice until further use. For cell disruption, cell pellets were kept on ice and resuspended in 10 ml of PBS supplemented with protease inhibitor (1/5 EDTA-free protease inhibitor cocktail tablet (Roche) in 10 ml 1x PBS). 1 ml of each cell suspension was transferred to a 2 ml reaction tube. After addition of glass beads (2 x 200 µl) cells were disrupted in a cryo mill at 30 s^-1^ for 10 min (4°C). Next, the suspension was centrifuged for 30 min at 16,200 x g(4°C). The cleared lysate (supernatant) was transferred to a new reaction tube and kept on ice. For the DNS assay, sucrose solution (25 mM sucrose in 1x PBS, pH 7.2) was mixed with 200 µl culture supernatant or 1:10 diluted (1x PBS) cleared cell extract. The mixture was incubated at 30°C and 600 rpm (stroke 3 mm). 60 µl samples were transferred to a new tube and incubated for 10 min at 95°C. Samples were taken after 0 h, 2 h, 6 h, 24 h, 30 h. Samples were stored at -20°C until determination of monosaccharide content using the DNS reagent. Samples were thawed at room temperature, 60 µl of DNS reagent (10 g/L 3,5-dinitrosalicylic acid (Acros Organics), 404 g/l potassium sodium tartrate tetrahydrate (Sigma-Aldrich), 0.4 M NaOH) is added. Reaction is accelerated by incubating at 99°C for 10 min. Samples were transferred to 96 well plate (Greiner, 96 flat bottom transparent) and absorbance at 540 nm was measured in microplate reader (Tecan Infinite 200 Pro). Values were normalized to the optical density for supernatant samples.

### 4.6 *Zea mays* seedling infection

Seeds of the *Zea mays* cultivar Early Golden Bantam were soaked in water overnight. 5 seeds were planted in soil and incubated in a temperature-controlled greenhouse (light and dark cycles of 14 hours at 28 °C and 10 hours at 20 °C, respectively). Seven-day-old seedlings (three leaf stage) were infected with indicated *U. maydis* strains. To this end, main cultures were inoculated with OD_600 nm_ of 0.25 and incubated at 28°C in a rotary shaker using baffled flasks. Cells were harvested upon reaching OD_600 nm_ 0.5 to 1 by centrifugation (1500xg 5 min, room temperature). The cell pellet was resuspended in sterile H_2_Obidest to an OD of 3.0 and the suspension was used for inoculating the seedlings at about 1 cm above soil level using 1 ml syringes with 0.4 x 13 mm needles (BD microlance 3). Approximately 200 µl cell suspension were used for each plant. After infection, plants were transferred to a CLF Plant Climatics plant chamber and incubated under the following conditions: 28°C, 18 h light (600 µEinstein) - 6 h dark cycles. Plants were watered with tap water every two days. Infection symptoms were scored after six and twelve days post infection (dpi) using established read-outs (15) including the categories “no symptoms”, “chlorosis”, “small tumors”, “large tumors”, “plant death”.

### 4.7 Recombinant production of invertases in *E. coli*

For biochemical characterization and Cryo-EM studies, recombinant *E. coli* BL21(DE3) cells harboring pET24d (C-6His) suc2_w/o_SP was harvested after 20 h cultivation, reaching a final OD_600_ of 11.5. *Escherichia coli* BL21(DE3) cells expressing recombinant Suc2^ΔSP^ from *U. maydis* were cultivated in 2× YT medium (tryptone, 16 g/L; yeast extract, 10 g/L; NaCl, 5 g/L; pH 7.0 ± 0.2) supplemented with 1% lactose for autoinduction and kanamycin (50 µg/mL). Cultures were incubated at 28°C with shaking at 180 rpm for 20 h. After cultivation, the final optical density at 600 nm (OD₆₀₀) was recorded, and cells were harvested by centrifugation. The cell pellet was resuspended in phosphate-buffered saline (PBS) and lysed on ice by sonication (30% amplitude, 30 s on/off intervals for 5 min). The soluble His-tagged protein with estimated extinction coefficient E_1%_= 17.18 was purified using immobilized metal ion affinity chromatography resulting in a volumetric yield of 55 mg/L. Next, overnight dialysis against 20 mM sodium acetate buffer, pH 5.5, was conducted. Protein concentration was determined by measuring absorbance at 280 nm, and the extinction coefficient was computed theoretically from the amino acid sequence (51).

For mass photometry heterologous expression of fungal invertases (tagged N-terminally with 6 x His tag) was performed in *E. coli* Rosetta pLysS according to Terfrüchte *et al.* (2017). Rosetta pLysS transformants harboring the respective expression plasmids were grown at 28 °C for 20 h in dYT supplement with kanamycin and 10 g/L lactose. The cells were harvested 4000 x g for 15 mins. The pellet was resuspended in Buffer A (20 mM HEPES pH 8.0, 20 mM KCl, 40 mM imidazole, 250 mM NaCl,) 10 mg/ml lysozyme solution was added. Cells were disrupted using an ultrasonic sonotrode (5 mm microtip (Heinemann, Portsmouth, NH, USA)) attached to the Cell Disruptor B15 device (Branson; stage 4, 3 × 30 s pulse, 5 repetitions). Cell lysates were subsequently centrifuged at 11.000 × g for 20 min (4 °C) and supernatants were subjected to IMAC purification following standard protocols. Elution was carried out stepwise with lysis buffer containing 250 mM and 500 mM imidazole, respectively. All obtained fractions were analyzed using SDS-PAGE and DNS assays were performed. Eluted proteins were stored at – 20 °C until further use. For mass photometry, samples were concentrated using centrifugal filters (Amicon Ultra 50k) centrifuged at 10,000 x g for 2 x 15 mins at to reach an end volume of 0.5 ml. Samples were then subjected to size-exclusion chromatography (SEC) using a Superdex S200 Increase 16/600 column. The peak fractions were analyzed using SDS- PAGE, for further use samples were pooled and concentrated again using centrifugal filters. Protein aliquots were stored at -80°C.

### 4.8 Mass photometry

Mass photometric measurements were performed using a TwoMP mass photometer (Refeyn Ltd, Oxford, UK) as described previously (52, 53), employing AcquireMP (Refeyn Ltd. v2.3) software for data acquisition. Mass photometry movies were recorded at 1 kHz, with exposure times varying between 0.6 and 0.9 ms, adjusted to maximize camera counts while avoiding saturation. Prior to use, microscope slides (70 x 26 mm) were cleaned for 5 min in 50% (v/v) isopropanol (HPLC grade in H_2_0_bidest_) and H_2_0_bidest_, followed by drying with a pressurized air stream. Silicon gaskets to hold the sample drops were cleaned identically and fixed to clean glass slides immediately before the measurement. The instrument was calibrated using the NativeMark Protein Standard (Thermo Fisher Scientific). The protein concentration during measurement of the different invertases was 30-70 nM. For each measurement, a fresh gasket well sas used. To identify the focus plane, 18 μL of fresh PBS buffer (RT) was added to a well, the focal position was determined and locked using the autofocus function. For each acquisition, 2 μL of diluted protein was added to the well and the liquid was thoroughly mixed. Three individual measurements were performed for each sample. DiscoverMP software was used for data analysis.

### 4.9 Recombinant protein production and purification

Heterologous expression of UmSuc2 (tagged N-terminally with 6 x His tag) was performed in *E. coli* Rosetta pLysS using plamid pET15b_His-Suc2^ΔSP^ (pUX492) according to Terfrüchte *et al.* (2017) (43). For synthesis of Suc2 Rosetta pLysS transformants harboring the expression plasmid were grown at 28 °C for 20 h in dYT supplement with kanamycin and 10 g/L lactose. The cells were harvested 4000 x g for 15 mins. The pellet was resuspended in Buffer A (20 mM HEPES pH 8.0, 20 mM KCl, 40 mM imidazole, 250 mM NaCl,) 10 mg/ml lysozyme solution was added. Cells were disrupted using an ultrasonic sonotrode (5 mm microtip (Heinemann, Portsmouth, NH, USA)) attached to the Cell Disruptor B15 device (Branson; stage 4, 3 × 30 s pulse, 5 repetitions). Cell lysates were subsequently centrifuged at 11.000 × g for 20 min (4 °C) and supernatants were subjected to IMAC purification following standard protocols. Elution was carried out stepwise with lysis buffer containing 250 mM and 500 mM imidazole, respectively. All obtained fractions were analyzed using SDS-PAGE and DNS assays were performed. Eluted proteins were stored at – 20 °C until further use.

### 4.10 Enzyme activity assay

Invertase activity was investigated using the saccharolytic 3,5-dinitrosalicylic acid (DNS) method to quantify the reducing sugars liberated from sucrose substrate. Standard reactions were performed in 50 mM acetate buffer (pH 5.5) at 30°C for 2 min in a final volume of 100 µL, using 0.01 µM purified enzyme. The reaction was terminated by the addition of equal volume of DNS reagent, followed by heating at 80°C for 5 min. Absorbance was measured at 540 nm, and the amount of reducing sugar produced was determined using a glucose calibration curve (22, 35). Negative controls (without enzyme) were included, all assays were performed in triplicate, and the results represent the mean of three experiments and error bars denote the standard deviation (SD). Kinetic parameters (K_m_, V_max_ and K_cat_) were determined by measuring reaction rates over a sucrose concentration range of 2-1000 mM under the standard assay conditions described above. Data were fitted to the Michaelis-Menten equation by non-linear regression. The turnover number (K_cat_) was calculated as K_cat_ = V_max_/[E], where [E] is the enzyme concentration, and is reported in s⁻¹.

### 4.11 Cryo-electron data collection and image processing

Cryo-electron microscopy (cryo-EM) data were acquired using a Talos Arctica transmission electron microscope (Thermo Fisher Scientific) operated at 200 kV and equipped with a K3 direct electron detector (Gatan). A total of 6,108 movie stacks were collected at a calibrated physical pixel size of 0.41 Å, with a total accumulated electron dose of approximately 50 e⁻/Å² and a nominal defocus range of –0.8 to –2.2 µm. Automated data collection was performed using EPU. Data processing was performed using CryoSPARC v4.7.1 (Figs. S4 and S5). The collected movie frames were aligned and dose-weighted using the Patch Motion Correction algorithm, followed by contrast transfer function (CTF) estimation with Patch CTF Estimation. Particles were initially auto-picked using the Blob Picker, extracted in 512 × 512 pixel boxes, and subsequently binned to 256 × 256 pixels for faster processing (Fig. S4). After multiple rounds of 2D classification (Fig. S5), approximately 1.8 million high-quality particles were selected and used to generate an initial three-dimensional model via ab initio reconstruction. Further heterogeneous refinement and non-uniform refinement steps were performed using 1.2 million particles. The final consensus map, refined with C1 symmetry, reached a resolution of 2.26 Å based on the gold-standard Fourier shell correlation (FSC) criterion of 0.143 (Fig. S5). Refinement imposing C2 symmetry yielded an improved nominal resolution of 2.19 Å.

### 4.12 Model building and refinement

The atomic model was initially built from the C1-symmetric cryo-EM map using ModelAngelo and subsequently refined by manual adjustments in Coot v1.1.15, including detailed fitting of the fructose density. The final real-space refinement was performed in Phenix v1.21.2, yielding a model comprising 1,071 protein residues and two fructose molecules, with no water molecules modeled. The refined structure displays excellent geometry (r.m.s.d. bond lengths 0.002 Å and bond angles 0.59°) and good stereochemistry (MolProbity score 2.10, clashscore 17.97, 96.2 % of residues in favored Ramachandran regions and 0.2 % outliers), and shows a strong overall fit to the cryo-EM density, with map–model correlation coefficients of 0.89 (mask) and 0.84 (volume) and a mean ligand CC of 0.70.

### 4.13 Microscopy and staining

Microscopy pictures were taken with an openFrame Microscopy System (Cairn GmbH, Ettlingen, Germany) equipped with the SCMOS sensor KINTEIX-22MM-M-C (Teledyne Photometrics, Tucson, AZ, USA) and an LDI-7 solid-state laser (89north, Williston, VT, USA) as illuminator. GFP imaging utilized 470 nm excitation and a 510 nm emission filter. Image acquisition was performed using Micro-manager (54). The microscopic analysis of yeast-like growing cells grown in CM-glucose was conducted at an OD_600_ of 0.5. For protoplast generation, a standard protocol was employed, using VinoZyme as a cell wall-degrading agent (45).

## 5 Data availability statement

Atomic coordinates of *U. maydis* Suc2^ΔSP^ have been deposited in the Protein Data Bank (PDB) under accession code 30SI (Extended PDB ID pdb_000030SI). The corresponding cryo-EM map, has been deposited in the Electron Microscopy Data Bank (EMDB) under accession code EMD-57998. The notebook for calculating growth rates is publicly available via https://github.com/Tanvir96rwth/Growth_rate_analysis.git.

## 6 Author contributions

T.B., A.K., H.Y. and J.C.H. designed and conducted the experiments. A.K. performed the cryo-EM sample preparation and data analysis, and contributed to the writing and preparation of the cryo-EM figures and results under the supervision of K.A.S.. T.H. generated the Jupiter notebook for growth rate analysis. F.A. generated structural predictions and structural conservation data. T.B., J.C.H. and K.S. designed and prepared the figures and tables. J.K., K.A.S., A.M. and K.S. supervised the project. F.A. and K.S. prepared the manuscript with input from all co-authors.

## Acknowledgements

We acknowledge Michael Feldbrügge for general support and constructive comments on the manuscript. Bettina Axler for excellent support in molecular cloning and strain generation. We also thank Lea Geissel and Hanna Göhlmann who contributed to strain generation and Sarah Goos for her valuable preliminary work on sucrose metabolism. We thank Gabriel Mendoza-Rojas for assistance with protein purification, Marius Benedens and Alexej Kedrov for mass photometry experiments and Johannes Postma for microscopy support. The authors gratefully acknowledge the electron microscopy imaging and access time granted by the life science EM facility of the Ernst-Ruska Centre at Forschungszentrum Jülich.

