## Supplementary Information for "Disentangling the sucrose metabolism of the corn smut *Ustilago maydis* reveals unexpected complexity"

**Running Header:** Alternative sucrose metabolism in *U. maydis*

#### Supporting Information

The Supporting Information contains Figures S1-S11 and Tables S1-S4.

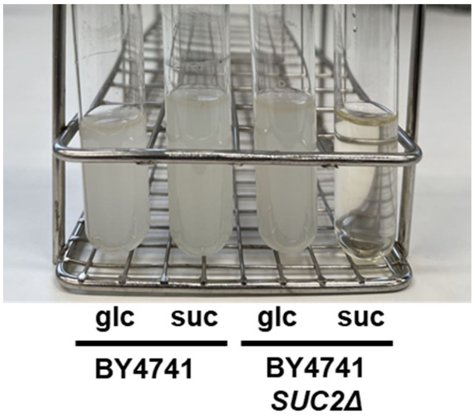


**Figure S1: Tester strain for invertase activity.** The picture shows cultivation tubes with indicated strains grown for 1 d on glucose (glc) or sucrose (suc) as single carbon source. In contrast to its progenitor BY4741 (MATa *his3Δ1 leu2Δ0 met15Δ0 ura3Δ0*), tester strain BY4741 *SUC2Δ* does not grow on sucrose.

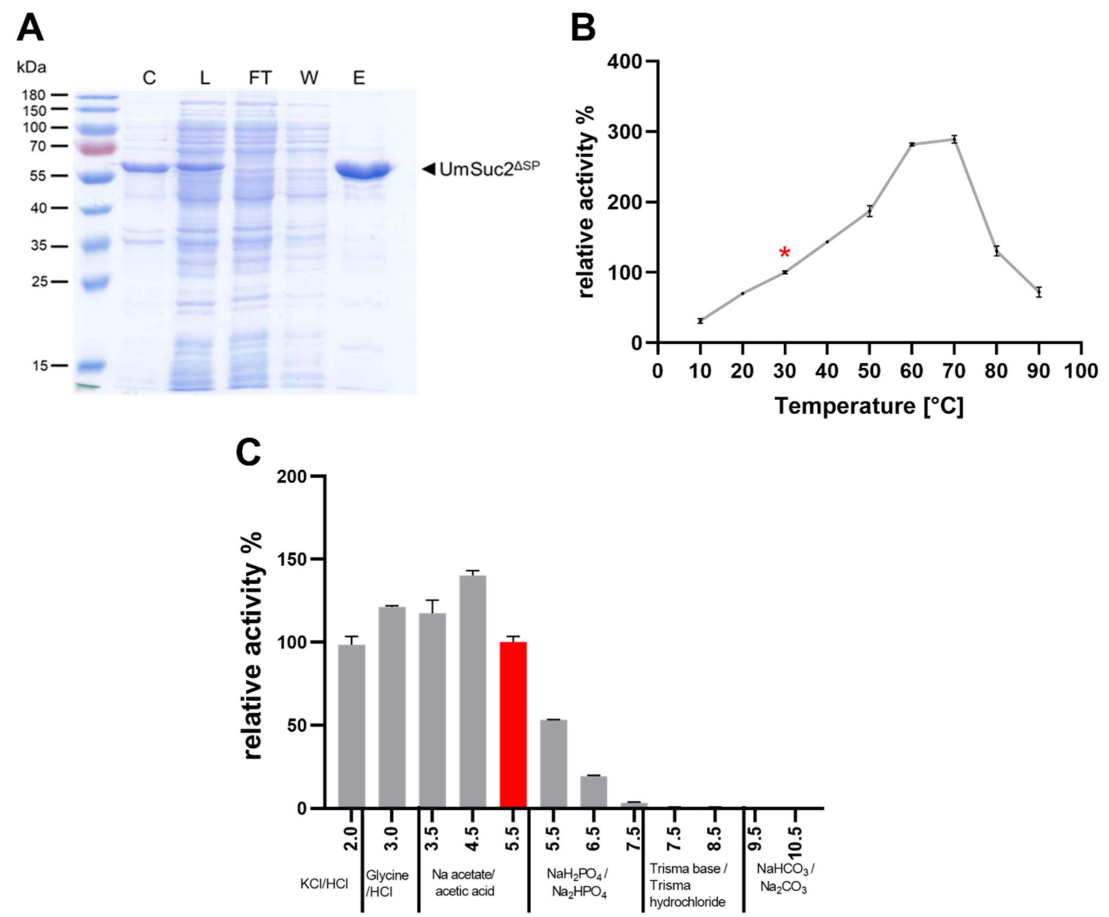


**Figure S2: Production and biochemical characterization of recombinant UmSuc2$^{\Delta SP}$. A,** Purification of UmSuc2$^{\Delta SP}$ produced in *E. coli* BL21 using the pET system. The protein carries a C-terminal His-tag used for purification via $Ni^{2+}$-chelate chromatography. Fractions are labelled as follows: C, cleared cell lysate; L, cell lysate; FT, flow-through; W, wash; E, elution. **B/C,** Effect of different temperatures (**B**), and pH (**C**) on the activity of recombinant UmSuc2$^{\Delta SP}$. Results are expressed relative to the control (marked in red or with red asterisk). Enzyme kinetics (Fig. S3) were determined at the parameters of the control. Error bars represent standard deviation of independent triplicates (control set to 100%).

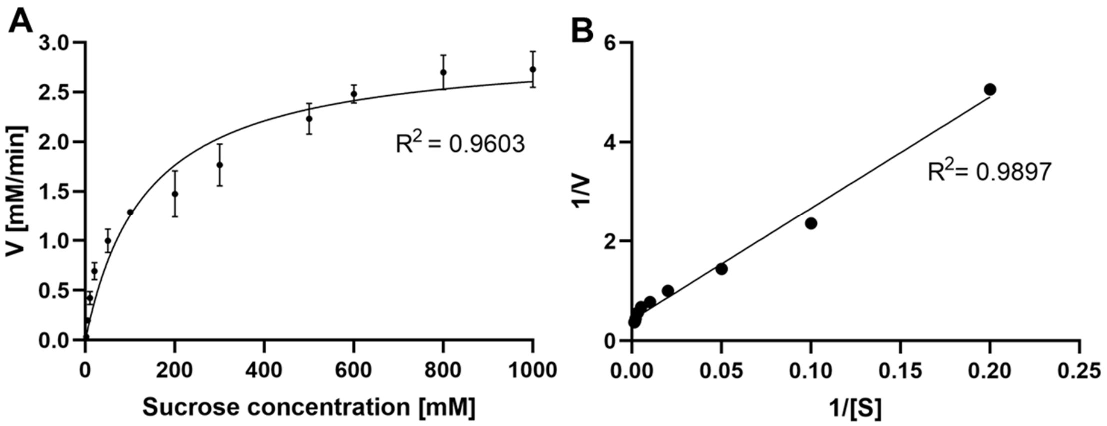


**Figure S3: Enzyme kinetics of UmSuc2.** Kinetics were determined at the parameters indicated in red in Fig. S2. **A**, Michaelis-Menten plot for determining the kinetic parameters of purified UmSuc2$^{\Delta SP}$ produced in *E. coli* BL21(DE3). Sucrose concentration was varied between 2 and 1000 mM. Values represent the mean of three determinations; bars represent the standard deviation. Equation: $y = 0.436\ln(x) - 0.5397$. **B**, Lineweaver-Burk plot, equation: $y = 22.42x + 0.4211$.

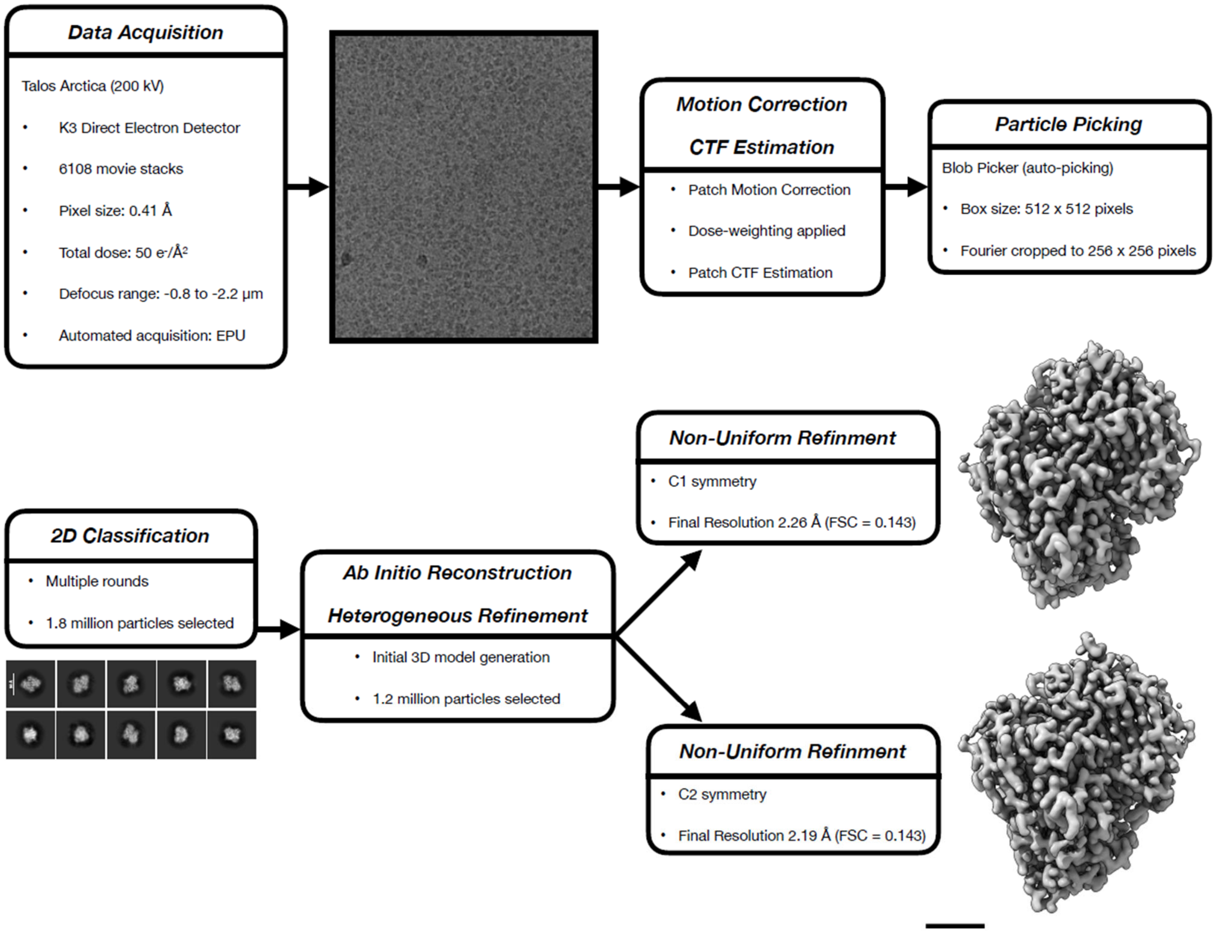


**Figure S4: Cryo-EM data acquisition and processing workflow.** Movies were collected on a Talos Arctica operated at 200 kV with a K3 direct electron detector, at a physical pixel size of 0.41 Å, a total dose of 50 $e^{-}/Å^{2}$, and a defocus range of −0.8 to −2.2 µm using automated acquisition in EPU. Movie frames were corrected for beam-induced motion and dose-weighted using Patch Motion Correction, and contrast transfer function parameters were estimated with Patch CTF in CryoSPARC. Particles were auto-picked with the Blob Picker, extracted in 512 × 512 pixel boxes, and Fourier cropped to 256 × 256 pixels prior to multiple rounds of 2D classification, yielding ~1.8 million high-quality particles for 3D analysis. An initial 3D model was generated *ab initio* and subjected to heterogeneous and non-uniform refinement, resulting in final reconstructions at 2.26 Å (C1 symmetry) and 2.19 Å (C2 symmetry) according to the gold-standard FSC 0.143 criterion.

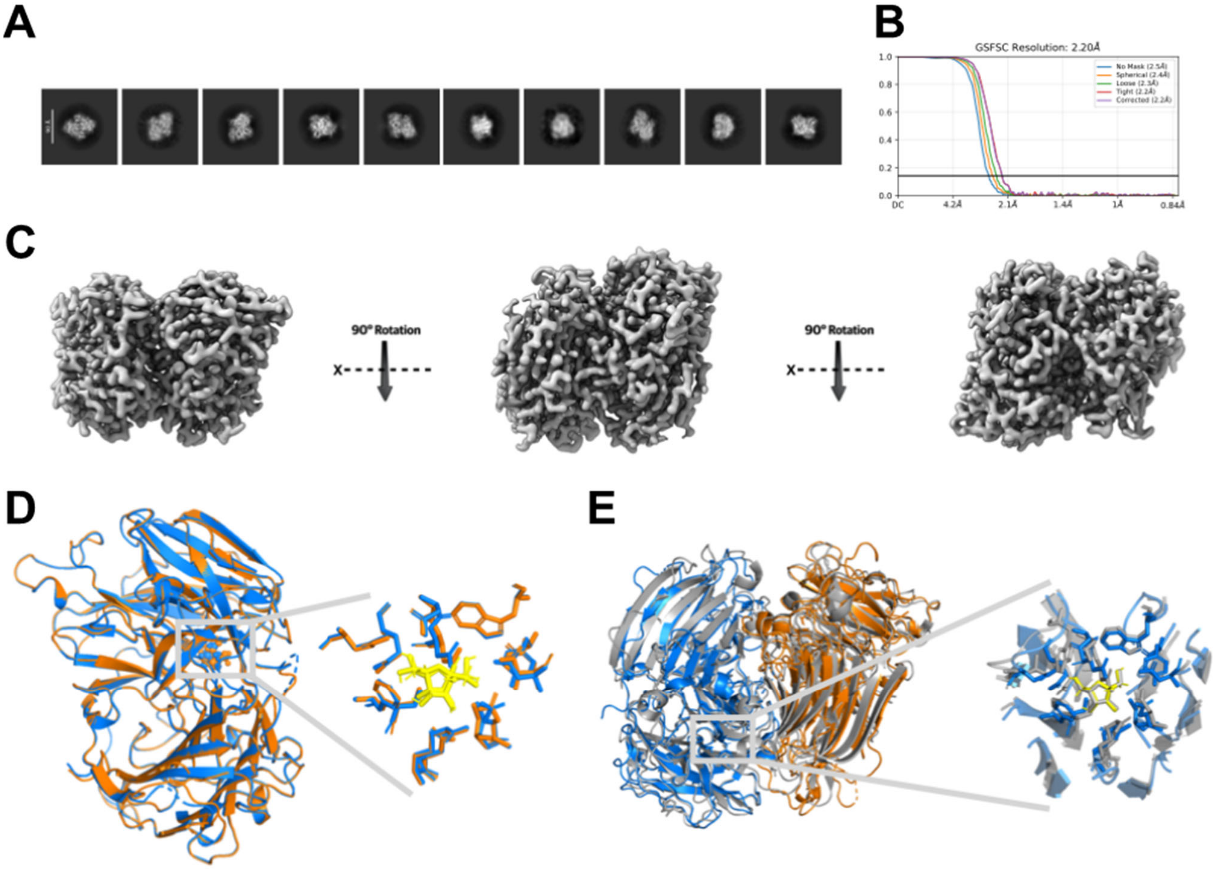


**Figure S5. Data processing, map quality, and structural comparison of recombinant UmSuc2$^{\Delta SP}$. A**, Representative 2D class averages obtained during cryo-EM data processing, illustrating a wide range of particle orientations used for 3D reconstruction. **B**, Gold-standard Fourier shell correlation (FSC) curve for the final reconstruction, indicating an overall resolution of 2.2 Å at the 0.143 criterion. **C**, Refined 3D reconstruction of dimeric invertase displayed in three orientations related by 90° rotations around the x-axis, highlighting the overall architecture and dimer interface. **D**, Superposition of the two monomers, with the dimer shown in two colors and a zoomed-in view of the active site **(**compare to Fig. 2C**)**; 3,397 atoms are aligned with a final RMSD of 0.194 Å, demonstrating that both protomers adopt virtually identical folds and active-site conformations. **E**, Structural comparison with the *S. occidentalis* invertase SoSuc2, showing superposition of the two enzymes and an enlarged view of the active site to illustrate conserved features and subtle differences in the substrate-binding environment.

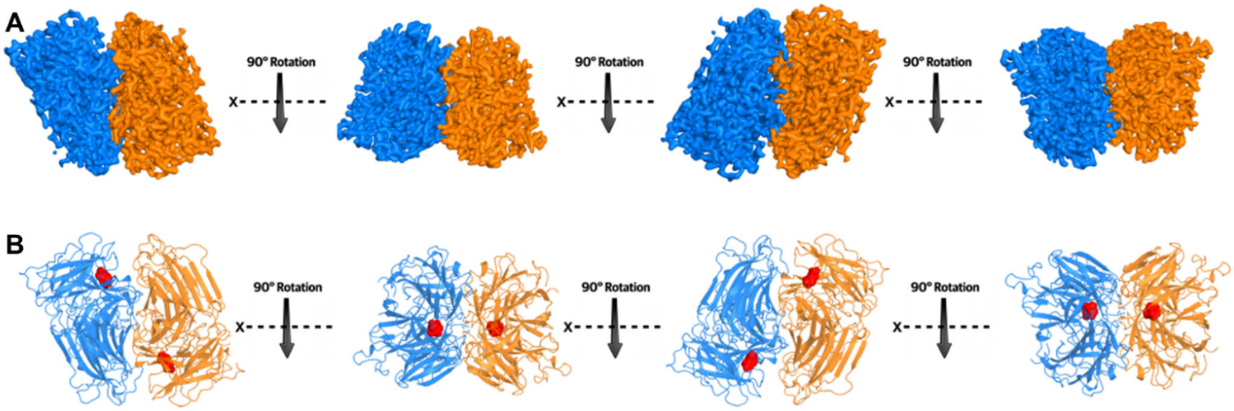


**Figure S6: Cryo-EM reconstruction of *U. maydis* Suc2**. **A**, Cryo-EM reconstruction of UmSuc2$^{\Delta SP}$ shown as a dimer, with the two protomers colored differently and displayed in four orientations related by 90° rotations around the x axis to illustrate the overall architecture and dimer interface. **B**, Corresponding atomic model of the dimer, colored as in panel A, with the fructose product density highlighted in red in both active sites to indicate the position of the catalytic pockets.

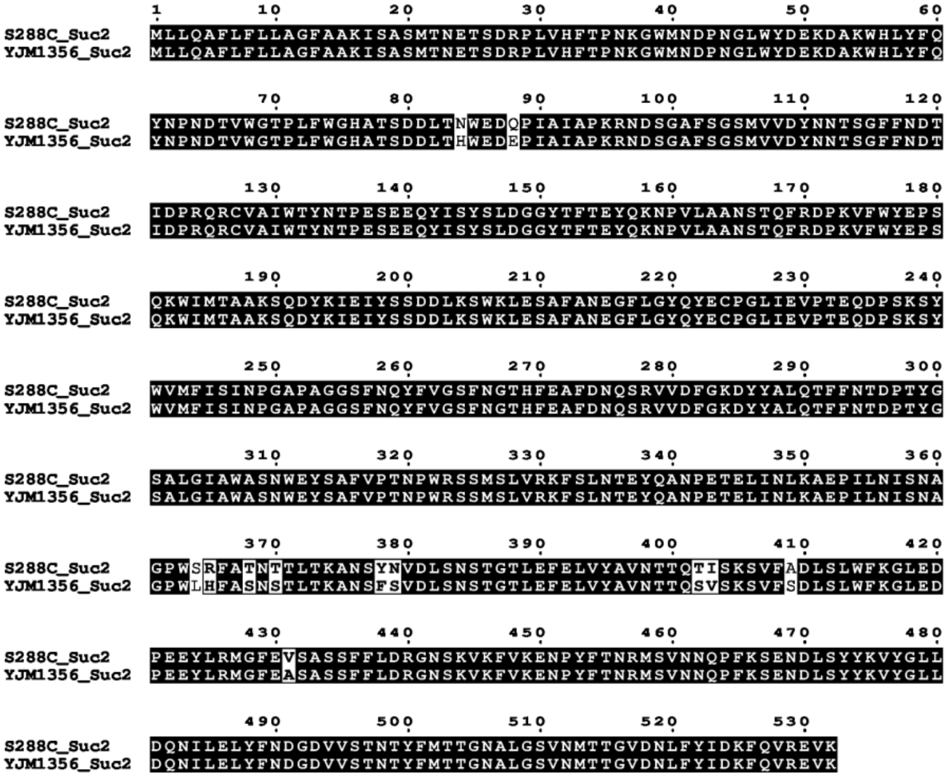


**Figure S7: Amino acid alignment of dimeric and higher-oligomeric ScSuc2.** Sequences of Suc2 versions originating from *S. cerevisiae* laboratory strain S288C (S288C_Suc2) and clinical isolate YJM1356 (YJM1356_Suc2) were compared. Identical residues are shaded in black while non-conserved residues have a white background. White boxes indicate similar amino acids. The alignment was generated with ClustalOmega and the program ESPript 3.0 (1).

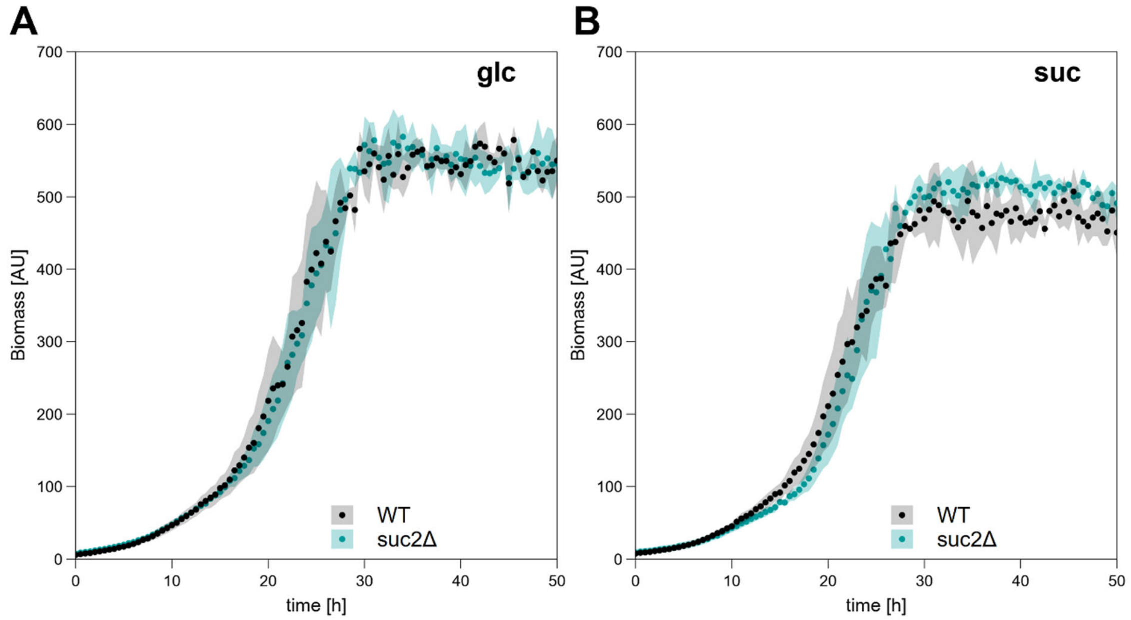


**Figure S8: A *suc2* deletion strain still grows on sucrose as single carbon source.** Growth of *U. maydis* laboratory strain AB33 (WT) and the indicated deletion mutant (suc2Δ) on glucose (**A**) and sucrose (**B**). The diagrams show backscatter data of indicated strains obtained in a micro-scale cultivation visualizing cell propagation. The assay was conducted in biological triplicates. Error bars depict standard deviation. AU, artificial units. glc, glucose; suc, sucrose. Note that the data of this figure is also included in Fig. 3B, C.

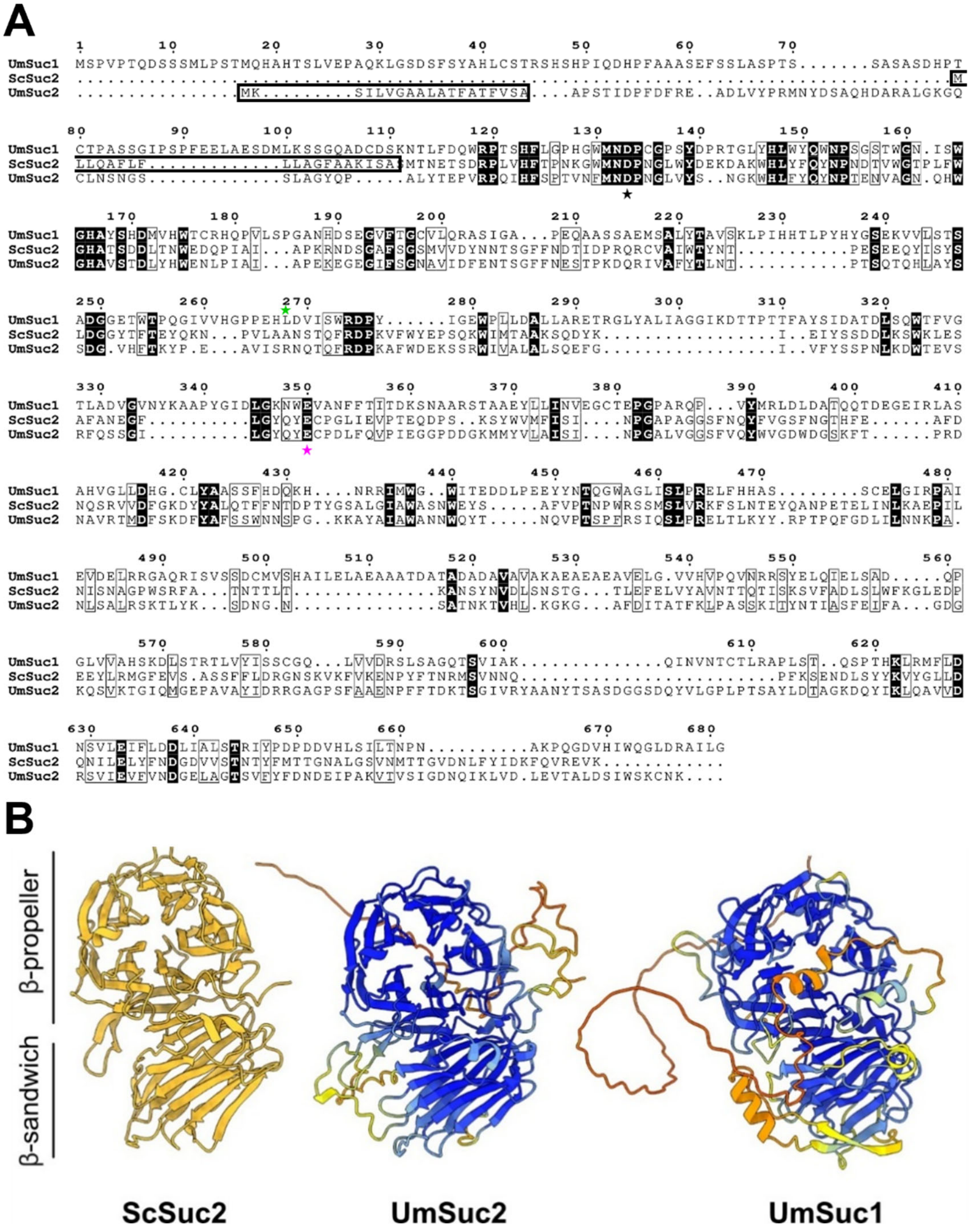


**Figure S9: Amino acid conservation and structure comparison of predicted *U. maydis* invertases. A,** Amino acid alignment of ScSuc2, UmSuc1 and UmSuc2 reveals low conservation on the secondary structure level while functional residues of the GH32 domain are conserved. Identical residues are shaded in black while non-conserved residues have a white background. White boxes indicate similar amino acids. Asterisks depict residues of the active side residues in the GH32 domain that are critical for invertase function based on homology to described ScSuc2 data (2). Black asterisk, nucleophile; pink asterisk, acid/base catalyst (2). The green asterisk indicates the position of the leucine 268 in UmSuc2 (compare main text). Predicted N-terminal signal peptides are indicated by open black boxes. The alignment was generated with ClustalOmega and the program ESPript 3.0 (1). **B,** Structural comparison using AlphaFold2 model of UmSuc1 to our structure of UmSuc2 and the published structure of a ScSuc2 monomer (PDB: 4EQV, yellow (3)).

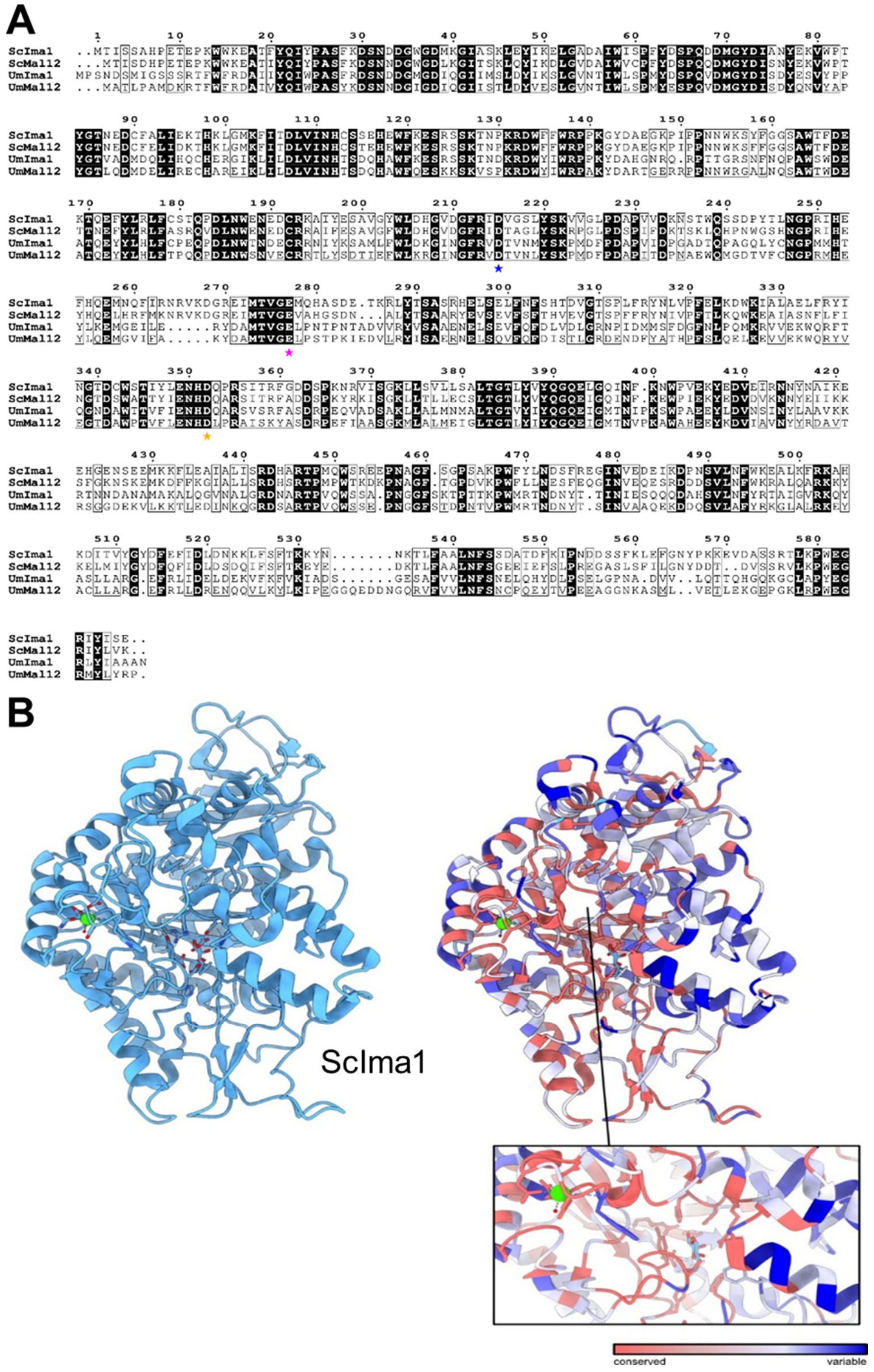


**Figure S10: Amino acid alignments, structural models and homologies of predicted (iso)maltases. A**, Amino acid alignment of ScIma1, ScMal12, UmIma1 and UmMal12. Identical residues are shaded in black while non-conserved residues have a white background. White boxes indicate similar amino acids. Blue asterisk depicts the nucleophile position, pink asterisk the proton donor residue, orange asterisk the transition state stabilizer annotated by sequence comparison (4). The alignment was generated with ClustalOmega and the program ESPript 3.0 (1). **B**, Overlay of AlphaFold2 predictions showing conservation of UmIma1 and UmMal12 as compared to yeast ScIma1 (cornflower blue, PDB-ID:3A4A). The right panel shows the sequence conservation mapped onto the ScIma1 structure with red regions indicating high conservation, while blue residues are less conserved.

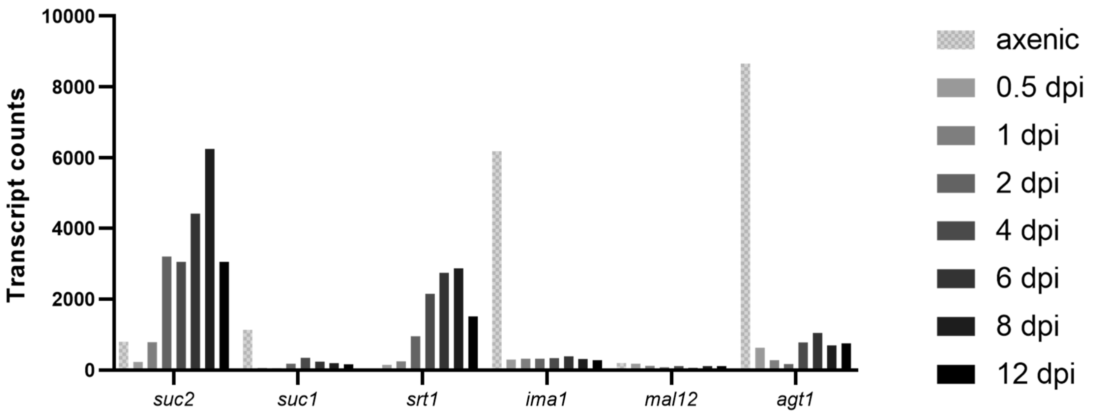


**Figure S11: Transcript abundance of candidate genes.** Expression data for indicated genes are derived from a published RNAseq approach (5). Yeast-like growing cultures (axenic) were grown on YEPSlight medium containing sucrose as major carbon source (axenic, hatched columns). Filled columns (shades of grey) indicate samples from infected plants (dpi, days post infection).

**Table S1**: Growth rates of indicated strains grown on 60 mM glucose or 30 mM sucrose as single carbon source. The growth rates were derived from data shown in Fig. 3D and 3E.

| Strain | Growth rate $\mu_{avg}$ on glucose [$h^{-1}$] | Growth rate $\mu_{avg}$ on sucrose [$h^{-1}$] |
|---|---|---|
| WT | 0.16 | 0.15 |
| Δsuc2 | 0.15 | 0.15 |
| Δsrt1 | 0.15 | 0.15 |
| Δsuc1Δsuc2 | 0.14 | 0.14 |
| Δsuc1Δsuc2Δsrt1 | 0.15 | 0.15 |
| Δsuc2Δsrt1 | 0.16 | 0.17 |

**Table S2**: Growth rates of indicated strains grown on 60 mM glucose, 30 mM sucrose or 30 mM maltose as single carbon source. The growth rates were derived from data shown in Fig. 4B-D.

| Strain | Growth rate $\mu_{avg}$ on glucose [$h^{-1}$] | Growth rate $\mu_{avg}$ on sucrose [$h^{-1}$] | Growth rate $\mu_{avg}$ on maltose [$h^{-1}$] |
|---|---|---|---|
| WT | 0.15 | 0.16 | 0.14 |
| Δsuc2Δsuc1Δsrt1 | 0.15 | 0.19 | 0.14 |
| Δsuc2Δsuc1Δsrt1Δima1 | 0.15 | 0.02 | 0.01 |
| Δsuc2Δsuc1Δsrt1Δima1Δmal12 | 0.15 | 0.01 | 0.02 |
| Δsuc2Δsrt1Δagt1 | 0.15 | 0.00 | 0.00 |
| Δsuc2Δagt1 | 0.15 | 0.00 | 0.00 |
| Δsuc2Δsuc1Δsrt1 Δima1Δmal12Δagt1 | 0.15 | 0.00 | 0.00 |

**Table S3.** Plasmids used in this study. Detailed information on the molecular cloning is available upon request. CbxR, genetic cassette mediating carboxin resistance; HygR, genetic cassette mediating hygromycin B resistance; NatR, genetic cassette mediating nourseothricin resistance; G418R, genetic cassette mediating geneticin resistance.

| **Plasmid (resistance)** | **Internal collection no.** | **Application** | **Reference** |
|---|---|---|---|
| pFLPexpC (CbxR) | pUMa1446 | FRT-mediated recycling of resistance cassettes | (6) |
| pSc_suc2D (HygR) | pUMa4567 | Deletion construct for *S. cerevisiae suc2* | This work. |
| pUm_srt1D (HygR, FRTm1) | pUX52 | Deletion construct for *U. maydis srt1* | This work. |
| pUm_suc1D (HygR, FRTm2) | pUX53 | Deletion construct for *U. maydis suc2* | This work. |
| pUm_suc2D (HygR, FRTm3) | pUX54 | Deletion construct for *U. maydis suc2* | This work. |
| pUm_agt1D (HygR, FRTwt) | pUX148 | Deletion construct for *U. maydis agt1* (HygR) | This work. |
| pUm_ima1D (HygR, FRTm5) | pUX149 | Deletion construct for *U. maydis ima1* | This work. |
| pSc_pTEF1:ima1Um_tENO2 (leu2) | pUX156 | Generation of complementation strain UX0119 | This work. |
| pSc_pPMA1:agt1_tADH1 (met15) | pUX159 | Generation of complementation strain UX0447 | This work. |
| pUm_agt1D (NatR, FRTwt) | pUX173 | Deletion construct for *U. maydis agt1* | This work. |
| pSc_pTEF1:mal12Um_tENO2 (leu2) | pUX179 | Generation of complementation strain UX0138 | This work. |
| pSc_pTEF1:suc1Um_tENO2 (leu2) | pUX180 | Generation of complementation strain UX0139 | This work. |
| pUm_mal12D (NatR, FRTm7) | pUX181 | Deletion construct for *U. maydis mal12* | This work. |

| | | | |
|---|---|---|---|
| pSc_pTEF1:suc2Um_tENO2 (leu2) | pUX197 | Generation of complementation strain UX0158 | This work. |
| pSc_pRPL18B:SUC2_ic_tENO2(LEU2) | pUX230 | Generation of complementation strain UX0230 | This work. |
| pUm_agt1D (NatR, FRTwt) | pUX476 | Deletion construct for U.m. agt1 (NatR)<br>Alternative to pUX173 with similar genetic gene deletion setup. | This work. |
| pUm_Psuc2:suc2-mKate2_Tnos (HygR, FRTm4) | pUX487 | *suc2::mKate2* fusion | This work. |
| pET24d (C-6His) suc2_w/o_SP | pUX492 | Heterologous expression of *U. maydis suc2* in *E. coli* (signal peptide deleted) | This work. |
| pET24d (6HN) suc2_Sc w/o SP | pUX804 | Heterologous expression of *S. cerevisiae* S288C *suc2* in *E. coli* (signal peptide deleted) | This work. |
| pET24d (6HN) suc2_Sc YJM1356 w/o SP | pUX806 | Heterologous expression of *S. cerevisiae* YJM1356 *suc2* in *E. coli* (signal peptide deleted) | This work. |
| pUM_Potef:suc2_ATG2 mut_gfp:Tnos (cbx) | pUL81 | Deletion of second ATG in *suc2*, fused with *gfp* | This work. |

### References (Supplemental material)
